# AI-guided screen identifies probucol-mediated mitophagy enhancement through modulation of lipid droplets

**DOI:** 10.1101/2022.05.18.492560

**Authors:** Natalia Moskal, Naomi P. Visanji, Olena Gorbenko, Vijay Narasimhan, Hannah Tyrrell, Jess Nash, Peter N. Lewis, G. Angus McQuibban

## Abstract

Failures in mitophagy, a process by which damaged mitochondria are cleared, results in neurodegeneration, while enhancing mitophagy promotes the survival of dopaminergic neurons. Using an artificial intelligence platform, we employed a natural language processing approach to evaluate the semantic similarity of candidate molecules to a set of well-established mitophagy enhancers. Top candidates were screened in a cell-based mitochondrial clearance assay. Probucol, a lipid-lowering drug, was validated across several orthogonal mitophagy assays. *In vivo*, probucol improved survival, locomotor function and dopaminergic neuron loss in zebrafish and fly models of mitochondrial damage. Probucol functioned independently of PINK1/Parkin but its effects on mitophagy and *in vivo* depended on ABCA1, which negatively regulated mitophagy following mitochondrial damage. Autophagosome and lysosomal markers were elevated by probucol treatment in addition to increased contact between lipid droplets and mitochondria. Conversely, lipid droplet expansion, which occurs following mitochondrial damage was suppressed by probucol and probucol-mediated mitophagy enhancement required lipid droplets. Probucol-mediated lipid droplet dynamics changes may prime the cell for a more efficient mitophagic response to mitochondrial damage.

## Introduction

Clearance of damaged mitochondria is important for the survival of dopaminergic neurons, the loss of which is responsible for the classical motor symptoms of Parkinson disease (PD). PD is a progressive neurodegenerative disease characterized by rigidity, akinesia and bradykinesia as well as wide ranging non-motor symptoms such as anxiety, depression, sleep disturbances and loss of smell^1^. Current treatment options address symptoms but fail to disrupt the progression of the disease, thus a disease-modifying treatment remains a major unmet need. Mitochondrial dysfunction is clearly implicated in PD pathogenesis, so enhancing the removal of damaged mitochondria (mitophagy) might have potential as a therapeutic intervention^1^.

Evidence for the importance of mitophagy in PD pathogenesis comes from both sporadic and genetic cases. Several disease-causing, loss-of-function mutations in genes encoding proteins which mediate mitophagy have been identified, including PINK1 and PRKN ^2,3^. Additionally, several disease-causing mutations in genes not directly associated with mitophagy, have secondary negative effects on mitochondrial health or on the retrograde transport of damaged mitochondria to axons, such as SNCA, GBA and LRRK2 ^4–6^. Besides genetic causes of PD, mitochondrial damage and mitophagy impairment have also been widely implicated in sporadic disease, including the inactivation of proteins which mediate mitophagy^7–9^. The clear connection between this pathway and the health of dopaminergic neurons emboldened us to search for novel mitophagy-enhancing compounds as potential disease-modifying therapeutics for PD.

A review by *Georgopoulos et al.*, describes several compounds currently known to stimulate and potentiate mitophagy^10^. However, most of these compounds also induce mitochondrial damage or apoptosis. While these are bona fide mitophagy enhancers, induction of mitochondrial damage or apoptosis would likely preclude their clinical use as this may worsen PD pathogenesis. To address the unmet need for mitophagy enhancers with therapeutic potential, we employed a computational approach using artificial intelligence to identify previously uncharacterized mitophagy enhancers. We have previously successfully used this strategy, that detects patterns and associations across several large datasets to identify compounds which are similar to a user-defined training set of positive controls, to identify drugs with disease modifying potential for PD that target aggregation of alpha synuclein^11–13^.

Here, using a similar approach, we screened a candidate list of molecules from the DrugBank database for similarity to known mitophagy enhancers. Many of the drugs have already been safely administered to humans. Repurposing drugs from other indications offers the opportunity to accelerate the clinical trials pipeline, given the presence of pre-existing pharmacological and toxicological information about candidate compounds^14^.

In a previous screen, we focused on compounds which accelerated the transition of the E3 ubiquitin ligase, Parkin, from the cytosol to the mitochondria-a step which is integral to this mitophagy pathway^15^. While this step is highly amenable to microscopy-based phenotypic screening, the limitations of this approach include possible omission of hits which function downstream of Parkin or that target other mitophagy pathways^16^. In this present screening iteration, we evaluated the clearance of damaged mitochondria from cells. Ultimately, if this downstream step is improved, then the negative consequences of mitochondrial damage in the dopaminergic neurons may be mitigated^17^. This effort led to the identification of a compound which enhances mitophagy following mitochondrial damage and led us to elucidate its mechanism of action, resulting in the identification of a newfound role for the ATP binding cassette transporter A1 (ABCA1) in mitophagy, through its effects on lipid droplets dynamics.

## Results

### Artificial intelligence (AI) simplifies mitochondrial clearance screen for mitophagy enhancers

We employed a computational approach using artificial intelligence (IBM Watson for Drug Discovery) to identify drugs amenable to repurposing as PD therapeutics. To identify candidate compounds, an *in silico* screen was performed to identify potential mitophagy enhancers from the DrugBank database (https://www.drugbank.ca), based on their similarity to positive control mitophagy enhancers. Positive controls were selected from a review article about pharmacological enhancers of mitophagy^10^. Ultimately, our training set comprised of the following 7 drugs: PTEN-induced putative kinase 1 (PINK1) activator kinetin; poly ADP-ribose polymerase (PARP) inhibitor olaparib, p53 inhibitor pifithrin-alpha; nicotinamide (NAD^+^ accumulation) and sirtuin1 activators resveratrol, fisetin and SRT1720.

First, we validated our model with a leave-one-out cross validation where each of the 7 training set compounds was excluded from the training set and re-ranked amongst the 3231 candidate drugs (Figure 1A). A retrospective analysis of Medline abstracts published up to and including 2014 was performed as further validation of the model. In this analysis, olaparib was omitted from the training set and placed in the candidate list for re-ranking. Using 194 available abstracts, the model ranked olaparib 126th amongst the candidate drugs (top 3.9%, Figure 1A).

**Figure 1.**
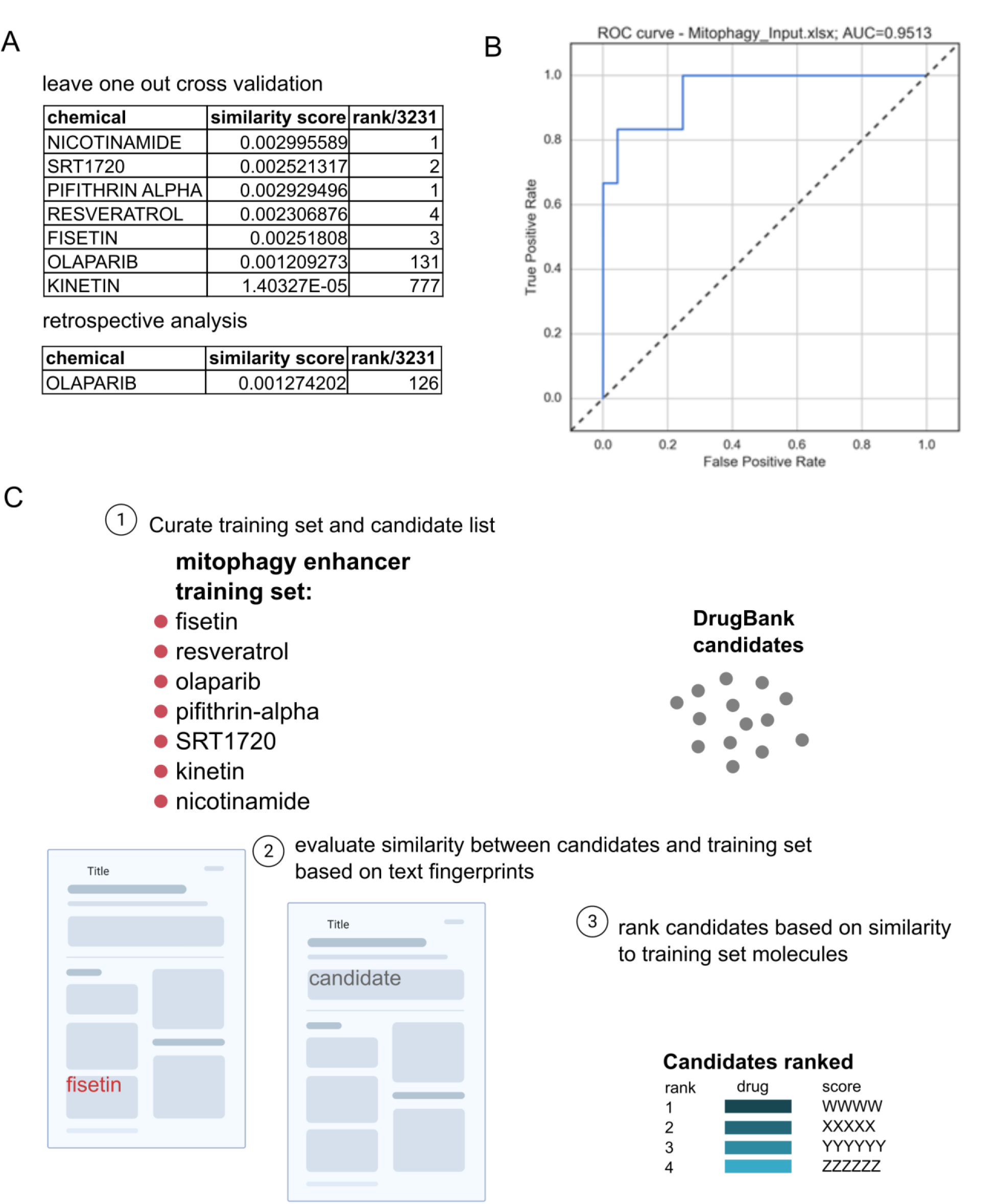
*in silico* screen to identify candidate mitophagy enhancers in DrugBank database of drugs amenable to repurposing (**A**) Leave one out cross validation and retrospective analyses were performed to evaluate the ability of the model to identify bona fide mitophagy enhancers. Similarity scores between each chemical assessed and the mitophagy enhancer training set were calculated and used to assign a rank out of 3231 DrugBank molecules from most to least similar in ascending order. (**B**) Leave-one-out cross validation results were used to construct a receiver operating characteristic curve which demonstrates the predictive performance of the model. The area under the curve for the ROC curve is 0.9513. (**C**) Following validation, the model was deployed to identify new mitophagy enhancers from the DrugBank candidates based on information from a wide array of sources including PubMed literature, patent filings and biological databases.

Taken together, this indicated that in the semantic model generated from Medline abstracts, the training compounds had strong predictive power over each other. A Receiver Operating Characteristics curve (ROC) was constructed using this data and an area under the curve (AUC) was calculated to assess the predictive power of the model (Figure 1B). An AUC value of 1 indicates perfect predictive ability, while <0.5 indicates predictive ability worse than random chance. The AUC for the model created by IBM Watson for Drug Discovery was 0.9513

Ultimately, we assessed 3231 candidate drugs from the DrugBank database for semantic similarity to the training drugs (Figure 1C). IBM Watson for Drug Discovery creates text fingerprints for all the training set molecules, in addition to the candidate molecules. The candidate molecules consist of FDA-approved small molecules, protein/peptide drugs and nutraceuticals. Many of the candidate molecules can be repurposed due to the lack of adverse effects associated with their administration in other disease contexts. The fingerprints created by IBM Watson for Drug Discovery encapsulate the words and phrases which are associated with a particular chemical entity in abstracts published on Medline. IBM Watson for Drug Discovery computed a similarity score for each candidate entity, and they were ranked from highest to lowest accordingly (Data S1). Due to overlap between compounds which cause apoptosis and/or mitochondrial damage and ones which induce mitophagy, two lists were cross-referenced to filter out the candidate molecules with any association to either the term “apoptosis” or “mitochondrial damage” (Data S2).

Next, the top 79 most similar candidate compounds identified by IBM Watson for Drug Discovery were screened in HeLa cells stably expressing GFP Parkin and mito-DsRed. Cells treated with the ionophore carbonyl cyanide m-chlorophenyl hydrazone (CCCP), which depolarizes the mitochondrial membrane potential, for 24 hours, undergo mitophagy resulting in loss of mito-DsRed signal in a large percentage of cells (Figure 2C, S1B)^18^. By quantifying the percentage of cells with low/no mito-DsRed signal, it is possible to assess mitochondrial turnover. Prior to CCCP treatment, the cells are pre-treated with 1 μM of the candidate small molecule (Figure S1A). In cells pre-treated with DMSO instead of small molecules, 78 ± 2.83% of cells have low/no mito-DsRed signal, which we henceforth refer to as mitochondrial clearance (Figure S1B). MAD z-scores were calculated, as described in prior mitophagy screens for each of the 79 compounds (Figure 2A, Data S3)^19^.

**Figure 2.**
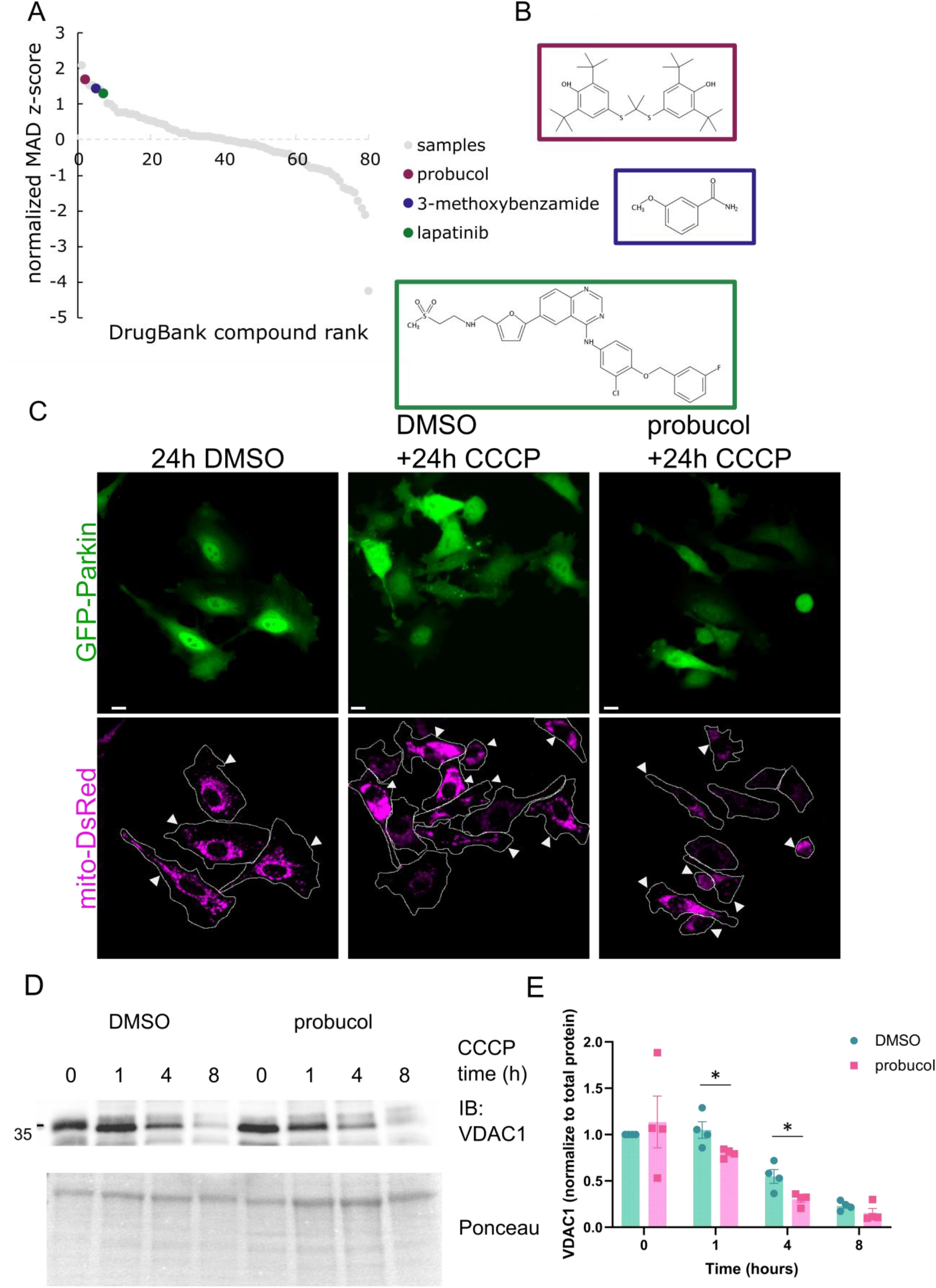
Cell-based mitochondrial clearance screen to evaluate candidates identified *in silico* **A**)Normalized MAD z-scores of 79 candidate DrugBank molecules screened in the mitochondrial clearance screen. Probucol, 3-methoxybenzamide and lapatinib are highlighted amongst other compounds in descending order of rank. **B**) Chemical structures of hit molecules highlighted in A). **C**) HeLa cells expressing GFP-Parkin and mito-DsRed. Cells were pre-treated with small molecules (1 μM) for 2 hours followed by 24-hour treatment with CCCP (10 μM) to induce prolonged mitophagy, resulting in loss of mito-DsRed signal from many cells (mitochondrial clearance). Arrows denote cells which retained mitochondrial signal. The percentage of remaining cells with no/low mito-DsRed signal was calculated as the screening readout. **D**) Immunoblotting using antibodies for outer mitochondrial membrane protein VDAC1 was performed on lysates from cells pre-treated with small molecule (1 μM) prior to CCCP (10 μM) time course. **E**) Quantification of VDAC1 levels normalized to Ponceau staining to assess protein loading. Data information: Normalized MAD z-score values were calculated based on two independent screening replicates in A). 4 independent biological replicates were performed for E). Bars represent mean values and error bars represent SEM. * indicates p-value <0.05. Statistical analysis was performed using an unpaired two-sided student’s t-test to compare DMSO and probucol at each time point.

Dichlorocopper ranked as 1/79 in our screen. Copper has known effects on mitophagy and general autophagy, more broadly^20,21^, so its recovery as a top hit was consistent with the goal of the screen. 3 compounds were selected for re-testing based on MAD z-score values and lack of evident toxicity in cells (Figure 2A), thereby producing a hit rate of 3.8%, which is higher than the hit rate in most high throughput screening campaigns which typically ranges between 0.01-0.14%^22^. Upon re-testing these compounds in an orthogonal mitochondrial clearance assay, probucol increased the extent to which mitochondrial substrates were degraded following induction of mitochondrial damage. Immunoblotting was performed to visualize levels of the inner mitochondrial membrane substrate ATP synthase F1 subunit alpha (ATP5A) (Figure S2) and the outer mitochondrial membrane protein voltage-dependent anion-selective channel protein 1 (VDAC1) following induction of mitochondrial damage with CCCP treatment.

Predictably, VDAC1 levels declined over the CCCP time course. While probucol treatment does not increase VDAC1 degradation under basal conditions, it promoted enhanced degradation of VDAC1 over the CCCP time course (Figure 2D, E).

### Probucol augments later stages of mitophagy

Upstream of the degradation of damaged mitochondria is the targeting of mitochondria to lysosomes. The mitoQC assay can be used in both cells and *in vivo* to probe this precise step in mitophagy. Briefly, cells were transfected with Cerulean-Parkin and RG-OMP25, a plasmid containing mCherry and GFP in tandem with the mitochondrial targeting sequence of OMP25 (Figure 3A). Upon localization to acidic lysosomes, GFP signal is quenched, which distinguished mitochondria localized to lysosomes because they appear as puncta with red-only signal. Less than 5 red-only puncta are present in cells under normal conditions, but the number of puncta increases following induction of mitochondrial damage^23^. Cells with greater than 5 red-only puncta corresponding to mito-lysosomes were classified as ‘mitophagic’ (Figure 3A). Following 6-hour CCCP treatment, 40.25 ± 6.3% cells are mitophagic compared to 13.32 ± 4.83% under normal conditions. A higher percentage of probucol-treated cells were mitophagic (66.29 ± 5.6%, Figure 3B), compared to all other treatments including with the other candidate compounds.

**Figure 3.**
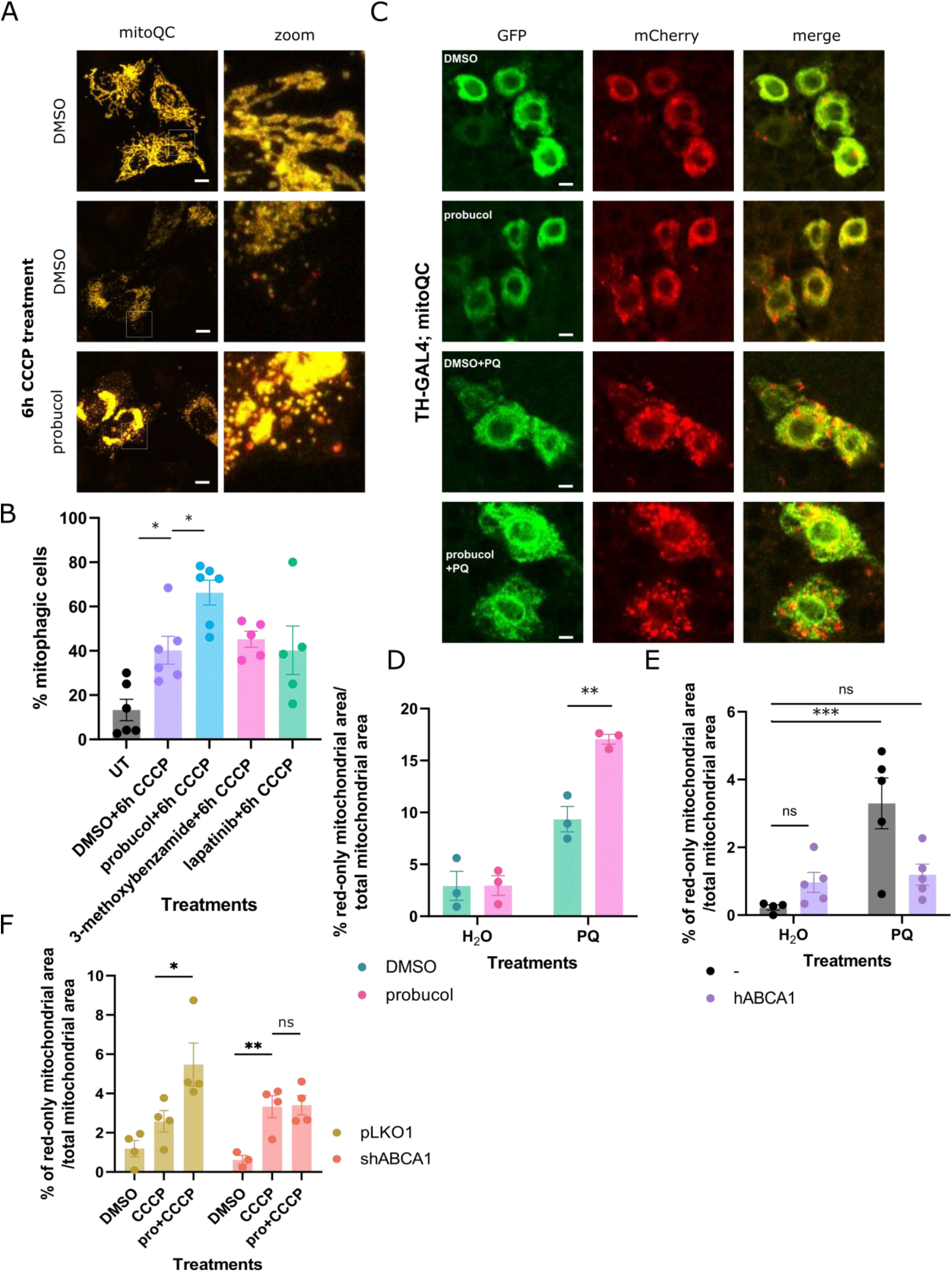
Probucol increased the targeting of mitochondria to lysosomes through ABCA1. **A**) The mitoQC reporter was expressed in HeLa cells along with Cerulean-Parkin. Mitophagy was stimulated by treating cells with CCCP for 6 hours along with either DMSO or candidate molecules from the screen. Mitochondria appear as red-only puncta when localized to acidic cellular compartments. **B**) Cells with >5 red-only puncta were defined as mitophagic and the mean percentage of mitophagic cells in each treatment condition is displayed. **C**) 7-day old flies expressing the mitoQC reporter in dopaminergic neurons were fed food supplemented with probucol in combination with either paraquat or water. **D**) Immunostaining using antibody against tyrosine hydroxylase (TH) segmented and defined dopaminergic neurons. The mean percentage of red-only mitochondrial area in each dopaminergic neuron is displayed. **E**) Control flies and flies co-expressing the human ABCA1 transgene and the mitoQC reporter were administered probucol in the presence and absence of paraquat. The mean percentage of red-only mitochondrial area in TH-positive neurons is displayed. See Appendix Figure S4D for corresponding images. **F**) Mitophagy was assessed in HeLa cells stably expressing mitoQC and Cerulean-Parkin which were transfected with either pLKO1 vector or with shABCA1. The mean percentage of red-only mitochondrial area per cell is displayed. See Appendix Figure S4C for corresponding images. Data information: Results are representative of at least 3 biological replicates, each indicated by data points in B, D, E and F. Bars represent mean values and error bars represent SEM. *, ** and *** indicative p-values <0.05, 0.01 and 0.005, respectively. At least 40 cells were assessed for B, D, E and F respectively. Dopaminergic neurons from at least 2 fly brains were analyzed for each treatment with at least 10 ROI per brain. Statistical analysis was performed using ANOVA and Dunnett’s multiple comparison correction.

To further substantiate these findings, the mitoQC reporter was expressed in the dopaminergic neurons of flies fed food supplemented with DMSO or probucol alone or in combination with paraquat, a mitochondrial toxin which causes PD in humans and PD-related phenotypes such as loss of dopaminergic neurons and locomotor impairments in model organisms such as flies^24^. Paraquat induces a mitophagy response characterized by an increase in mito-lysosomes in dopaminergic neurons^25^. Mitophagy further increased in flies co-administered probucol along with paraquat (Figure 3C, D). Basal mitophagy did not differ in the dopaminergic neurons of flies fed either DMSO- or probucol-supplemented food.

To determine whether probucol’s canonical target, ABCA1, is involved in its effects on mitophagy, we performed several genetic manipulations to change ABCA1 levels in cell culture or in the dopaminergic neurons of flies. The same mitophagy assay used to characterize probucol’s effects was also performed in this context.

Human ABCA1 and mitoQC transgenes were co-expressed or mitoQC was expressed alone in the dopaminergic neurons of flies (Figure 3E, S4D). Once again, paraquat treatment increased the percentage of red-only mitochondrial puncta, which represent mitochondria localized to lysosomes (Figure 3E). Basal mitophagy was unaffected by overexpression of the transgene. However, in flies fed paraquat, ABCA1 reduced the percentage of mitochondrial area localized to lysosomes (Figure 3E). Immunoblotting was performed to confirm expression of human ABCA1 (Figure S4A).

We also reduced ABCA1 levels with shRNA (Figure S4B) in cells expressing mitoQC and Cerulean-Parkin and treated cells as in Figure 3A. While probucol increased the percentage of red-only mitochondrial area indicative of mito-lysosomes in cells transfected with the control pLKO1 vector, probucol no longer increased mitophagy in cells transfected with shABCA1 (Figure 3F).

Follow up experiments probed steps further upstream in the mitophagy cascade, including PINK1-mediated phosphorylation of ubiquitin at S65 and recruitment of Parkin to damaged mitochondria (Figure S3). Phosphorylation of mitochondrial ubiquitin S65 following mitochondrial damage did not further increase following probucol treatment based on immunostaining and immunoblotting experiments (Figure S3A, B and D). Parkin recruitment to damaged mitochondria likewise was not increased by probucol treatment (Figure S3C), indicating that this mitophagy enhancer likely exerts its effect on steps further downstream of Parkin recruitment or through other Parkin-independent mitophagy pathways^16^.

### Probucol improves mitochondrial damage-induced phenotypes across several animal models of PD

*Drosophila* and *Danio rerio* serve as PD model systems, as they replicate much of human PD pathogenesis and display phenotypes which reflect human disease presentation. Flies and zebrafish exhibit loss of dopaminergic neurons resulting in impaired locomotion following exposure to PD-causing toxins such as 1-methyl-4-phenylpyridinium (MPP^+^) and paraquat ^24,26^. After replicating these features in the two model organisms, we tested probucol’s disease-modifying potential. The climbing ability and survival of flies declined following paraquat administration (Figure 4A, B) but co-treatment with probucol improved locomotor function and survival (Figure 4A, B).

**Figure 4:**
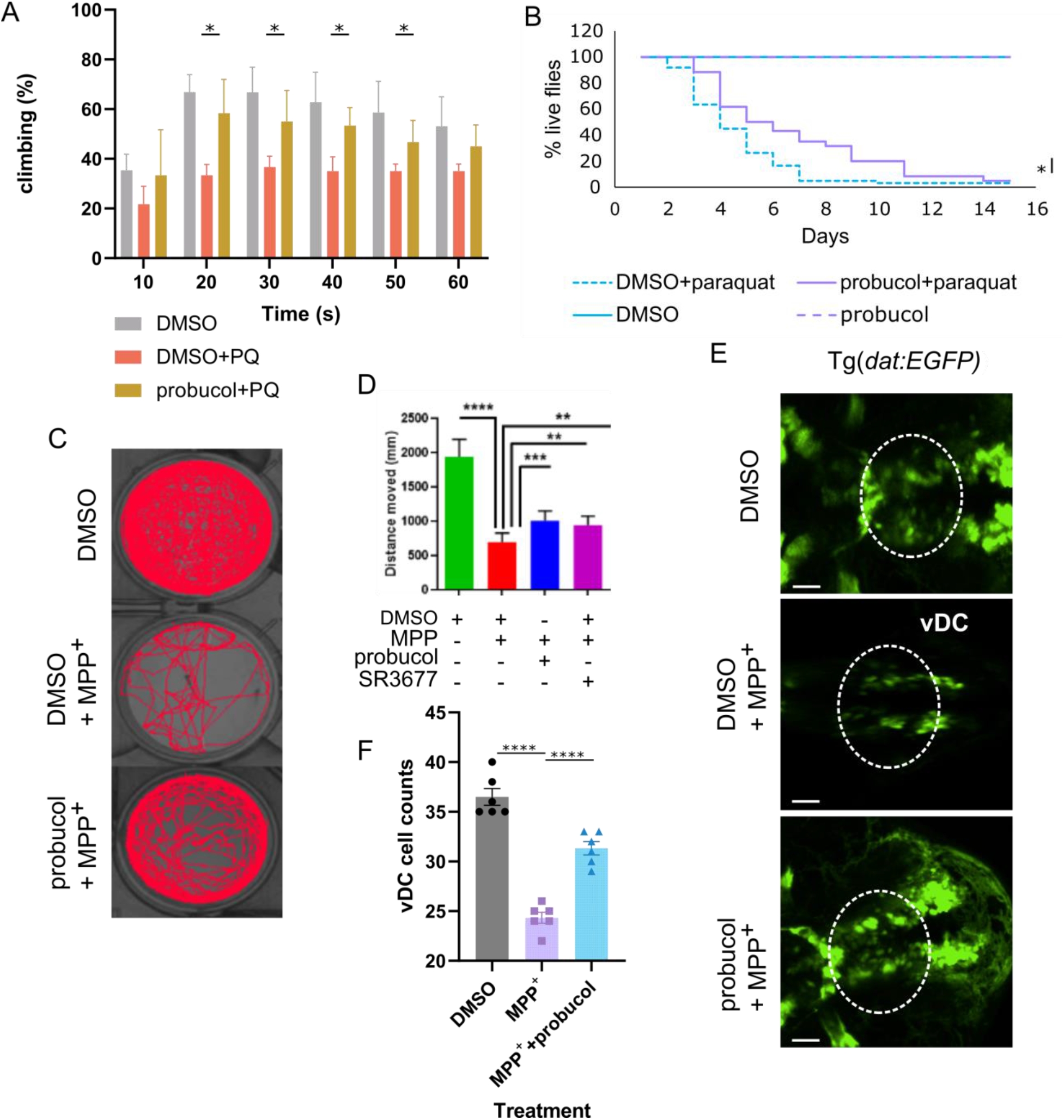
Probucol alleviated PD-related phenotypes which arose from mitochondrial dysfunction *in vivo*. **A**) Effect of probucol administration on paraquat-induced climbing defect. The percentage of flies to climb across a height of 12.5 cm. **B**) Effect of probucol administration on survival, in the presence and absence of paraquat co-treatments. **C**) Single particle tracking traces to visualize distance travelled by zebrafish in wells following administration of DMSO, probucol and MPP^+^, as indicated. **D**) Distance travelled by zebrafish in each treatment group in C, in addition to following treatment with positive control compound SR3677. **E**) Tg*(dat:EGFP)* zebrafish brains were imaged following treatment with DMSO, probucol and MPP^+^. Dashed lines highlight the vDC region of interest. **F**) The number of vDC neurons in each treatment group in E was quantified. Data information: At least 3 independent biological replicates were performed for each experiment, with individual replicates depicted by data points in F. Bars and error bars represent mean and SEM values, respectively. Unpaired student’s t-tests were performed to analyze data in A, log-rank tests were used to analyze survival data and one-way ANOVA analysis was performed to analyze data in D and F along with Dunnett’s multiple comparison correction. *, **, *** and **** indicate p-values <0.05, <0.01, <0.005 and <0.001, respectively.

Zebrafish embryos were incubated in MPP^+^ in combination with probucol or DMSO. Following 6 days of incubation, the movement of adult zebrafish was captured using ZebraBox. The distance travelled by the zebrafish exposed to DMSO in combination with MPP^+^ was visibly reduced compared to zebrafish incubated in DMSO alone (Figure 4C, D) but, the addition of probucol to the MPP^+^ increased the distance travelled by the zebrafish (Figure 4C, D). SR3677, a chemical inhibitor of Rho-associated protein kinase 2 (ROCK2) was employed as a positive control. We previously characterized this compound as a mitophagy enhancer and found that it improved locomotor decline in flies fed paraquat^25^.

In addition to toxin-based models of PD, we also tested probucol’s effect in a genetic model of mitochondrial dysfunction. Specifically, in heteroplasmic flies with approximately 90% of mitochondrial DNA (mtDNA) that contained a temperature-sensitive mutation in mitochondrial cytochrome c oxidase subunit I (*mt:ColI^T300I^*). Shifting these flies to a non-permissive temperature causes mitochondrial dysfunction resulting in climbing defects and significant reduction of lifespan^25,27,28^. Climbing and lifespan were improved in heteroplasmic *mt:ColI^T300I^* flies fed food containing probucol rather than DMSO (Figure S6). Much like paraquat, the phenotypes displayed by these flies arise from mitochondrial dysfunction and probucol reduced their severity, possibly by promoting mitophagy in this context as well, given that mitophagy has previously been shown to remove mitochondria bearing deleterious mutations^29^.

To directly probe the cell type impacted in PD, *Tg(dat:EGFP)* zebrafish embryos were co-incubated with MPP^+^ and probucol or vehicle control. Dopaminergic neurons can easily be visualized and counted in this transgenic model^30^. Following 1 day of incubation in MPP^+^, the number of dopaminergic neurons in the ventral diencephalon (vDC), which is analogous to the human nigrostriatal region, was reduced. In contrast, probucol-fed zebrafish retained more of their dopaminergic vDC neurons (Figure 4E, F). This reduced loss of dopaminergic neurons likely led to the probucol-mediated improvements to PD-related phenotypes.

### ABCA transporter and its effects on lipid droplets mediate probucol’s mitophagy enhancement

The extent of crosstalk between lipid homeostasis and autophagy reported in other studies and ABCA1’s role in lipid efflux led us to test whether probucol altered lipid droplets (LD). LDs have been found to form in response to starvation-induced autophagy and deferiprone-induced mitophagy^31,32^. LDs similarly increased significantly in cells treated with CCCP for 24 hours (Figure 5C, S5). Surprisingly, probucol treatment reduced the LD area expansion that occurred following mitophagy induction but had no effect on LD area under basal conditions (Figure 5C, S5). Lipid droplet area also increased in the dopaminergic neurons when flies consumed paraquat in their food. Probucol supplementation decreased the lipid droplet area expansion to levels comparable to those under basal conditions (Figure 5A, B).

**Figure 5:**
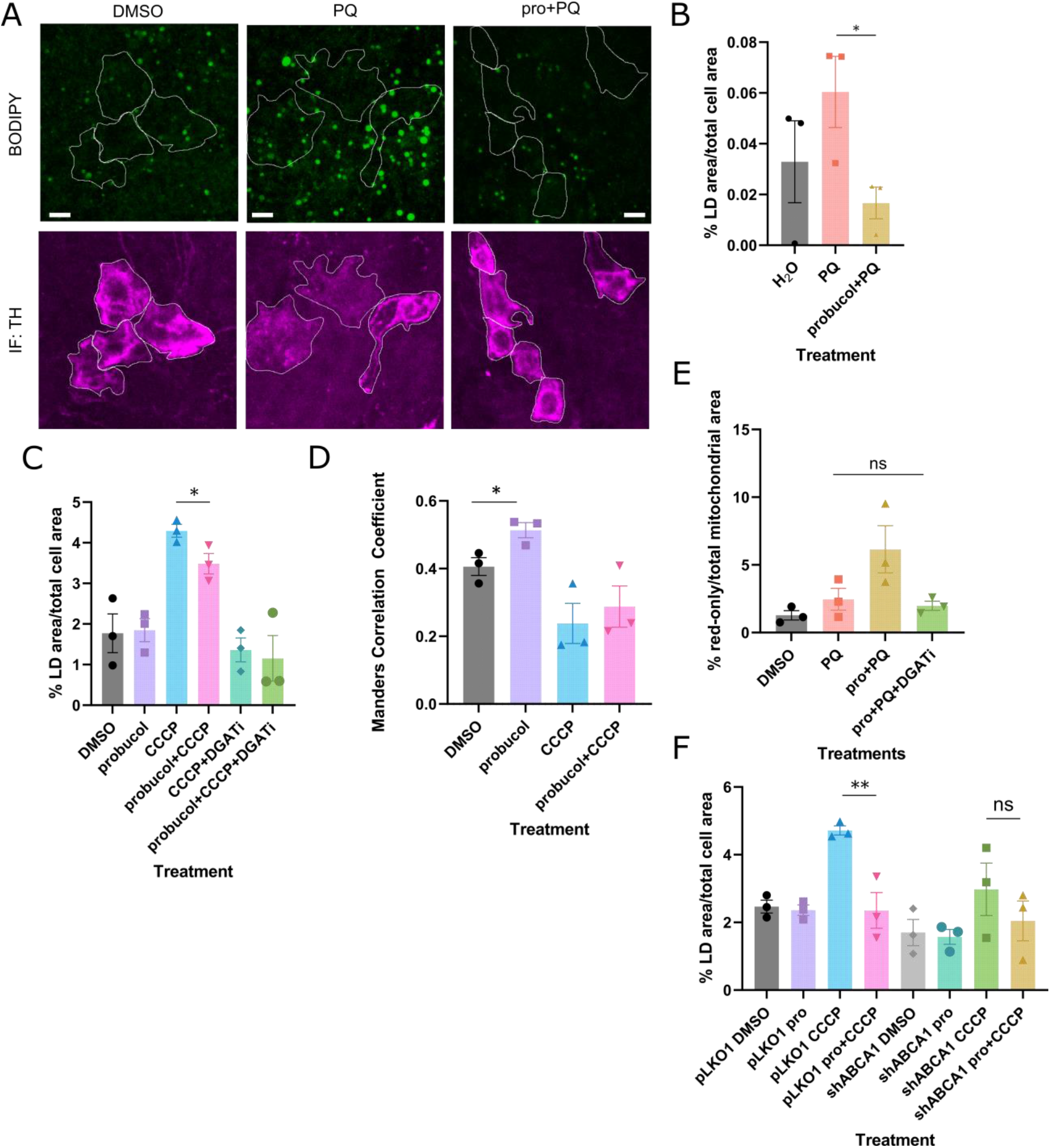
ABCA-mediated effects of probucol on mitophagy depended on lipid droplets, which increased proximity to mitochondria upon probucol treatment **A**) TH-positive dopaminergic neurons were segmented in BODIPY-stained brains from flies fed food supplemented with the indicated combinations of probucol and paraquat. **B**) The percentage of LD area over the total TH-positive cell area is quantified from A. **C**) The percentage of LD area over total cell area in HeLa cells treated with either DMSO or probucol alone or in the presence of CCCP with and without DGAT inhibitors. **D**) The overlap between BODIPY-stained LDs and mitochondria was assessed using the Manders Correlation Coefficient in HeLa cells treated as described in C). **E**) Probucol treatment was co-administered along with paraquat in the presence and absence of DGAT inhibitors. The percentage of red-only mitochondrial area was measured. **F**) HeLa cells were transfected with pLKO1 and shABCA1 plasmids and treated with the indicated combination of probucol and CCCP for 24 hours. BODIPY staining was used to quantify the percentage of cell area occupied by LDs. Data information: Results are representative of at least 3 independent biological replicates, as indicated by the data points. Mean values are displayed with error bars which represent the SEM. * and *** represent p-values <0.05 and 0.005, respectively. Statistical analysis was performed using ANOVA analysis, with multiple comparison correction for E and F. Unpaired student’s t-test analysis was performed to compare DMSO and probucol groups in B, C and D.

LD-mitochondria contacts have been observed in several studies. Peridroplet mitochondria localized to contact sites exhibit altered bioenergetic properties and serve as a source of ATP for LD expansion^33^. However, under conditions of high autophagic flux, LDs are thought to interact with mitochondria to buffer the lipotoxic species that are produced as byproducts of autophagy^31^. Interestingly, a greater proportion of lipid droplets colocalized with mitochondria in probucol-treated cells under basal conditions where mitochondria appeared intact and visibly elongated (Figure 5D, S5). Under the conditions in which this increased contact is observed, mitophagic flux is not elevated (Figure 3C, D), so it is unlikely that the contacts facilitate buffering of lipotoxic byproducts like in starvation-induced autophagy^31^.

Finally, to evaluate the importance of lipid droplets on the mitophagy enhancement conferred by probucol treatment, diacylglycerol acyltransferases (DGAT)1 and DGAT2 inhibitor treatment was added to suppress the lipid droplet expansion which occurred following prolonged mitochondrial stress (Figure 5C, S5). The addition of DGAT inhibitors to probucol treatment in the context of paraquat-induced mitochondrial dysfunction in flies attenuated probucol’s enhancement of mitophagy in the dopaminergic neurons of flies (Figure 5E). Mitophagy was no longer elevated by probucol in the context of paraquat, when DGAT inhibitors are present. Since DGAT inhibitors reduced LD area and attenuated probucol’s effect on mitophagy, this suggests probucol’s effect on LDs may be responsible for its subsequent effects on mitophagy.

Since probucol no longer exerted effects on mitophagy without lipid droplets and when ABCA levels are reduced, we probed whether ABCA1 affected probucol-mediated LD expansion. This critical step in probucol’s mechanism of action on mitophagy was no longer evident when ABCA1 levels were reduced with shRNA (Figure 5F, S4A).

### Probucol increases LC3 lipidation and lysosome abundance

Since early mitophagy steps were unaltered by probucol, downstream steps were assessed next. Immunoblotting can be used to evaluate the lipidation status of LC3. The lower molecular weight, lipidated form of LC3 (LC3-II) correlates with autophagosome levels and therefore increases as autophagy proceeds. However, it is important to note that increased LC3-II levels alone, in the absence of co-treatments with lysosome inhibitors, cannot directly indicate increased autophagic flux^34^. LC3-II levels increased following probucol treatment under basal conditions (Figure S7) in HEK293 cells with endogenous Parkin present at low levels. LC3-II levels were predictably higher upon mitophagy induction with CCCP treatment compared to basal conditions, as other studies show. However, the difference in LC3-II between DMSO- and probucol-treated cells did not persist under these conditions.

A recent report found that autophagy must be tuned to provide sufficient dynamic range to resolve differences in LC3 lipidation following manipulations such as drug treatments^35^. This can be accomplished by employing bafilomycin, an inhibitor of lysosome-autophagosome fusion at low doses so the effects of manipulations would be apparent. Experiments were repeated in HeLa cells which lack endogenous Parkin with the addition of low dose bafilomycin treatment. While the difference between DMSO and probucol treated groups under basal conditions was not as robust as in HEK293 cells, the addition of bafilomycin revealed a difference in LC3 lipidation between DMSO and probucol-treated cells in the CCCP group (Figure 6A, B).

**Figure 6:**
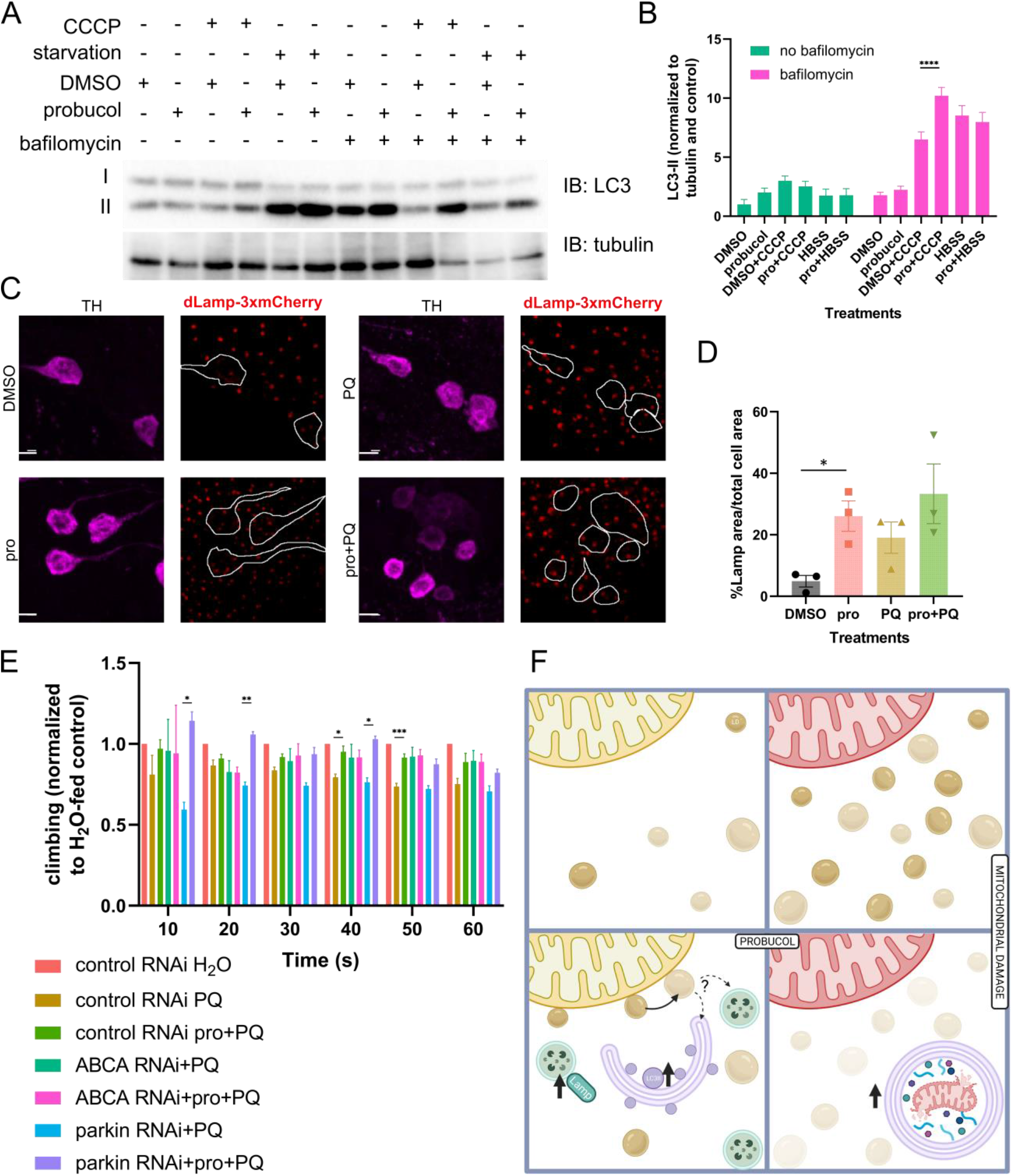
LC3 lipidation and lysosome area increased following probucol treatment and ABCA was necessary for probucol-mediated climbing improvements. **A**) HeLa cells were either incubated in DMEM, DMEM with CCCP or HBSS media for 6 hours in the presence or absence of bafilomycin. Lysates were separated by SDS-PAGE and immunoblotting was performed using antibodies which recognize LC3 and tubulin, as a loading control. **B**) Densitometry was performed to measure the levels of lipidated LC3-II, which were normalized to the tubulin loading control. **C**) Endogenously tagged *Lamp*-3xmCherry flies were fed food supplemented with DMSO or probucol in the presence and absence of paraquat. **D**) The Lamp area in each cell was measured as a percentage of total cell area, as defined by segmentation of TH-positive dopaminergic neurons. **E**) RNAi targeting either *parkin* or *ABCA* was expressed in the dopaminergic neurons with the TH-Gal4 driver. The effect of probucol on paraquat-induced climbing impairment was assessed in flies from the various genotypes. Climbing as a percentage of flies to cross 12.5 cm height is displayed. **F**) Hypothetical mechanism of mitophagy enhancement by probucol may involve upregulation of Lamp positive late endosomes/lysosomes and LC3-II positive mature autophagosomes, possibly arising from mobilization of lipid droplets adjacent to mitochondria. Lipid droplet expansion occurs following mitochondrial damage, but in the presence of probucol the abundance of lipid droplets is reduced, resulting in increased abundance of mito-lysosomes. Data information: Bars represent mean values and error bars correspond to SEM from at least 3 independent biological replicates, which are indicated with data points in graphs B and D. Statistical analysis to assess differences between DMSO and probucol within the same treatment condition or genotype group was performed using one-way ANOVA tests with multiple comparison correction in B and E, while unpaired t-tests were used to compare DMSO and probucol in D. *, * and *** indicate p-value <0.05, <0.01 and <0.005, respectively.

An endogenously mCherry-tagged *Lamp* reporter fly line was employed to assess lysosomes *in vivo*. Lamp is a lysosomal protein, which increases in abundance following paraquat treatment^36^. Dopaminergic neurons were segmented with TH immunostaining and the area occupied by Lamp-positive lysosomes within dopaminergic neurons was measured. As expected, paraquat addition to fly food increased abundance of Lamp-positive puncta (Figure 6C). Probucol treatment increased the area of lysosomes in the dopaminergic neurons, under basal conditions where no exogenous stimulus was added to trigger lysosome accumulation (Figure 6C, D).

To determine whether probucol’s effect on mitophagy were responsible for its ability to improve paraquat-induced climbing impairment, climbing assays were performed as described in Figure 4A, but the dopaminergic neuron-specific TH-GAL4 driver was used to drive expression of RNAi targeting candidate genes. In this manner, we dissected which factors were dispensable for probucol-mediated climbing improvements. Consistently with experiments in cells that showed that probucol had no effect on Parkin subcellular distribution (Figure S3), *parkin* RNAi did not abrogate the effects of probucol-mediated improvement of climbing impairment, which was evident in both the mCherry^RNAi^ and *parkin*^RNAi^-expressing flies (Figure 6E).

However, probucol treatment no longer improved climbing when *ABCA* RNAi was expressed in dopaminergic neurons (Figure 6E). In sum, these findings show that probucol treatment increased downstream autophagy steps such as autophagosome and lysosome biogenesis, seemingly priming the cells for a more rapid and efficient mitophagy response when damage strikes. ABCA, probucol’s target likely facilitates this effect through alterations to LD dynamics.

## Discussion

Using our screening approach, several previously characterized mitophagy enhancers were recovered both *in silico* and in cells. *In silico* screening identified staurosporine amongst the top hits (17/3231, top 0.5%), which is a well-characterized activator of mitophagy, but it was excluded from further experiments as it induces apoptosis^37^.

In our cell-based screen, dichlorocopper ranked as 1/79 in our screen. Copper binds to and increases the kinase activity of autophagy regulatory kinases ULK1 and ULK2^21^. ULK1/2 mediate autophagosome formation downstream of PINK1/Parkin^38,39^. Dichlorocopper would not have been identified had our mitophagy screen focused on Parkin recruitment, but nevertheless enhanced mitophagy, so the design of this screen may be superior to our previous screening approach^25^. The recovery of compounds with established mitophagy-promoting effects gave us confidence in the predictive power and efficacy of our dual screen. By filtering out compounds which induce mitochondrial damage or apoptosis, our effort was focused on identifying new compounds and mechanisms leading to mitophagy enhancement.

Interestingly, the set of compounds used to train our *in silico* model to identify mitophagy enhancers largely consisted of SIRT1 agonists (Figure 1C). SIRT1 affects mitophagy by upregulating the mitophagy receptor BNIP3 in aged mouse kidney^40^. Several compounds function by increasing the cellular NAD^+^ pool, which is a cofactor for SIRT1. This bias led us to speculate that our dual screen would identify several more mitophagy enhancers that function through this common mechanism shared amongst the training set. Despite the bias, only 1 of the 3 final hits (3-methoxybenzamide) was a SIRT1 agonist. Interestingly, a recent phase I clinical trial has demonstrated efficacy for nicotinamide, an NAD^+^ precursor, in PD, so the mechanism of action encompassed in the training set is likely nevertheless a relevant disease target^41^.

Ultimately, the screen identified probucol, a drug used to treat hypercholesterolemia prior to the advent of statins. The target which probucol inhibits is the ATP-binding cassette transporter ABCA1 and diverse assays in cells and flies which involved genetically reducing ABCA levels abolished probucol’s effects on mitophagy in cells and climbing in flies, while expressing human ABCA1 reduced mitophagy in dopaminergic neurons^42^. Experiments probing distinct steps in the mitophagy pathway found that probucol impacted mitophagy and *in vivo* phenotypes independent of PINK1/Parkin but required LDs, as pharmacological inhibition of LD biosynthesis abrogated probucol’s mitophagy enhancing effect.

Under normal conditions, probucol had several relevant effects: 1) it increased LD-mitochondria contacts, 2) increased late endosomes/lysosomes and 3) increased autophagosome lipidation. Importantly, probucol did not increase mitolysosome abundance under normal conditions but does so following mitochondrial damage. LDs adjacent to mitochondria can supply fatty acids during nutrient stress^43^ and can interact with and transfer lipids and proteins to both lysosomes and autophagosomes^44,45^. LD mobilization by lipase PNPLA5 is required to facilitate the formation of autophagic membranes, including in the context of mitochondrial damage^46^. Given that we observed increased abundance of late endosomes/lysosomes and mature autophagosomes under basal conditions, we speculate that the latter is true and may occur adjacent to mitochondria under basal conditions, but further investigation is required. Ultimately, the increased abundance of two components which can subsequently fuse to form mito-lysosomes when mitophagy is induced likely primes the cell for a more efficient and protective degradative response.

Cells and flies were subjected to prolonged mitochondrial damage in the form of CCCP treatment or paraquat feeding. Under these treatment conditions, we observed an increase in lipid droplet abundance. This is consistent with studies by *Nguyen et al.* and by *Long et al.* which demonstrate increase lipid droplet levels upon starvation-induced autophagy and deferiprone-induced mitophagy^31,32^. The two studies attribute different roles for LDs in these contexts. LDs buffer lipotoxic species which are generated as a byproduct of autophagic degradation and facilitate the subcellular transition endolysosomes undergo from the peripheries of the cell towards damaged perinuclear mitochondria, respectively. We did not investigate the reason for lipid droplet expansion occurs upon mitochondrial damage, but we did find that lipid droplets were necessary for probucol’s effects on mitophagy.

Lysosome position is critical for effective macroautophagy and found to be disrupted by inhibition of LD biosynthesis in deferiprone-induced mitophagy^32,47^. The increased abundance of LDs at mitochondria may facilitate the positioning of lysosomes away from the peripheries where they become active^47,48^. *In vivo*, clear overlap between lipid droplets and lysosomes was visible both in dopaminergic neurons and in other cells of the fly brain, supporting this possibility.

Interestingly, lipid droplet expansion following mitochondrial damage was reduced in probucol-treated cells and flies. This feature of probucol’s mechanism may be particularly relevant, given recent studies which found increased lipid droplet accumulation in the dopaminergic neurons of PD patients^49^. Likewise, the reduced accumulation supports the idea that LDs may be mobilized by probucol treatment to facilitate mitophagy^46^. Probucol mitigated both lipid droplet accumulation and mitochondrial damage-two features of PD pathogenesis. Probucol’s canonical target, ABCA1, is required for both probucol’s effects on mitophagy and on LD expansion. Targeting a point of crosstalk between these two pathogenesis pathways may be advantageous.

ABCA1 R219K gene polymorphisms impacts PD progression, as measured using the Hoehn and Yahr scale^50^. The K allele, which is associated with slower PD progression, also affects the lipid profile of those who carry the genotype. Compared to individuals carrying ABCA1 R219K RR or RK, ABCA1 R219K K genotype carriers have elevated high density lipoprotein cholesterol and lower triglyceride levels^51^. It remains to be determined whether the differences in lipid profiles are responsible for clinical differences between these groups.

Across toxin-based and genetic models of mitochondrial damage and PD in two different species, probucol improved survival, locomotor function and reduced the loss of dopaminergic neurons. A prior paper also found improvements to phenotypes caused by mitochondrial dysfunction in *C. elegans*. In this study, how probucol facilitates these improvements was not examined^52^. Given probucol’s promising effects in several preclinical animal models, it might be fruitful to mine human epidemiologic data for any associations between probucol treatment and reduced risk of PD, since it is still in use in Japan and China. Statins also increase mitophagy, but in a Parkin-dependent manner, unlike probucol^53^. Whether statins impact lipid droplet expansion following mitochondrial damage may represent an interesting avenue for future inquiry.

In conclusion, our study showcased a dual *in silico*/cell-based screening methodology which identified known and new mechanisms leading to mitophagy enhancement. ABCA1, which localizes to endolysosomes in cells, and regulates lipid homeostasis may serve as a mediator of crosstalk between lipid droplet dynamics and mitophagy since lipid droplets are required for the mitophagy enhancement conferred by probucol^54^.

## Methods

### Cells and tissue culture

All cell lines and sources are compiled in the Materials table. Cells were cultured in Dulbecco’s Modified Eagle Media supplemented with 10% fetal bovine serum. Cells were routinely tested for mycoplasma using the e-myco VALID mycoplasma testing kit (FroggaBio, 25239). Cells were maintained at 37°C temperature and 5% CO_2_ in a humidified atmosphere.

### Immunoblotting

Lysates were harvested with lysis buffer (0.1M Tris HCl, 0.01% SDS, pH 9) containing protease inhibitor cocktail (BioShop, PIC002.1) followed by 20 minutes of boiling and vortexing at 95°C. Lysates were then pelleted by high-speed centrifugation for 20 minutes at 4°C and supernatant was transferred into a new microcentrifuge tube for BCA assays to standardize protein loading across each experiment (Pierce, 23227).

Immunoblotting for VDAC1, ATP5A, ABCA1 was performed with 10% SDS-PAGE gels to separate proteins followed by transfer onto PVDF membrane (Immobilon, IPVH00010) at 110V for 80 minutes or 8-hour 38V transfer in the cold room. Samples to be probed for LC3 were separated on 15% SDS-PAGE gels instead. The transfer apparatus set up included an ice pack and a stir bar.

After Ponceau staining and imaging to assess overall protein loading, membranes were washed with TBST and blocked in 5% skim milk diluted in TBST for 30 minutes at room temperature. Incubation in primary antibodies diluted in 2.5% skim milk was performed overnight at 4°C. 3 5-minute TBST washes were performed prior to incubation of blots in secondary antibody diluted in 2.5% milk for 2 hours. 3 final 10-minute TBST washes were followed by visualization of proteins with ECL (BioRad, 11705062). Densitometry analysis was performed using ImageLab 6.0 software (BioRad) and the protein of interest was normalized to either Ponceau staining or loading controls such as GAPDH or tubulin. Antibodies used in this study are compiled in Materials table.

### Immunofluorescence

Cells grown on coverslips (1.5H thickness) were fixed with 4% PFA for 15 minutes, followed by 3 PBS washes. Permeabilization was performed with 0.1% Tx-100 incubation for 15 minutes, followed by 3 more PBS washes. Blocking with 10% goat serum diluted in PBS was performed for 30 minutes at room temperature, or overnight at 4°C. Coverslips were then incubated in primary antibody diluted in 1% goat serum (1:500) overnight at 4°C. The next day, following 3 PBS washes, coverslips were incubated in secondary antibody diluted in 1% goat serum (1:500) for 1-2 hours at room temperature. 3 10-minute PBS washes were followed by mounting onto slides with Fluoromount containing DAPI (Invitrogen 00495952).

Immunofluorescence of fly brains began with 20-minute fixation with 4% PFA, followed by 3x washes with 0.1% PBST (Tween-20 diluted in PBS) washes. Permeabilization was performed with 1% PBST (Triton-X-100 diluted in PBS) for 1 hour, followed by 3x washes in PBS. Blocking of fly brains in 10% goat serum for 1 hour was followed with overnight incubation at 4°C in primary antibody diluted in 1% goat serum. The next day, 10 minute 0.1% PBST (Tween-20 diluted in PBS) washes were followed by incubation in secondary antibody diluted in 1% goat serum for 2 hours. 3 more 10 minute 0.1% PBST washes and a final 10-minute PBS wash were performed prior to overnight incubation of fly brains in Fluoromount (Invitrogen 00495802) and mounting the next day onto slides. Coverslips (1.5H thickness) were sealed to the slide with clear nail polish. Antibodies used in this study are compiled in Materials table.

### Fly husbandry

Fly lines used in this study are compiled in Materials table*. Drosophila* were maintained at 25°C and at 70% relative humidity in 12-hour light/dark cycles and were fed standard yeast-molasses-sugar-agar formula. In the case of drug treatments, low melt agar fly food formula was composed as previously described^55^. Probucol was added to low melt agar fly food at 250 μM concentration and the equivalent volume of DMSO was used in control vials. DMSO concentration never exceeded 1% in experiments. Fly stocks used in the study and sources are listed in Materials Table.

### *In silico* screen for mitophagy enhancers in Drugbank

We have previously described the natural language processing methodology employed by IBM Watson for Drug discovery predictive analytics^11,12^. In brief, a set of candidate drugs were ranked according to semantic similarity to a training set of drugs known to have the desirable biological effect using natural language processing applied to published abstracts obtained from Medline.

#### Training set

A training set consisting of compounds listed in Figure 1A was composed with reference to a review about pharmacological modulators of mitophagy^2^. This list was curated to filter out any that were associated with mitochondrial damage or apoptosis (Data S2).

Compounds associated with key words such as apoptosis, depolarization and mitochondrial damage were excluded. The result was a set of 7 compounds with proven ability to induce mitophagy without an association with mitochondrial damage or apoptosis; Olaparib (PARP inhibitor), nicotinamide (NAD^+^ accumulation), pifitherin-α (p53 inhibitor), kinetin (PINK1 neo-substrate) and the SIRT1 activators resveratrol, fisetin and SRT1720.

#### Candidate set

3231 final candidates were filtered from the entire DrugBank database (https://www.drugbank.ca). Candidates with less than 5 published abstracts were removed.

#### Model Validation

1. **Leave one out cross validation:** Leave one out cross validation was performed. The ranking was run 7 times, with each training drug in turn removed from the training set and ranked among the other 3231 candidate drugs. Receiver Operating Characteristics curves were generated across the range of possible ranks to assess the model’s performance in a binary classification task. Data were analyzed in Python using the scikit-learn library.
2. **Retrospective analysis:** Olaparib was first published as having a mitophagy inducing effect in 2015^56^. To further validate our methodology, a ranking was performed restricted to abstracts published up to and including 2014. Olaparib was omitted from the known set and placed in the candidate list.

#### Ranking of candidate set and selection of candidates for validation

The final ranking was applied resulting in a list of 3231 candidate drugs rank ordered according to semantic similarity to the training set.

Following post-hoc analysis, the top 79 molecules ranked as bearing highest semantic similarity to the training set molecules, were purchased from Sigma. The identities of the compounds in our custom library are described in Data S3. These molecules were arrayed in a 96-well format at 10 mM concentration in DMSO.

### Screening

20 000 HeLa cells stably expressing mito-DsRed and GFP Parkin were seeded into clear, flat bottom, black polystyrene 96-well plates (Corning, CLS3603) and incubated at 37°C with 5% CO2 overnight. A final concentration of 1 μM of the compound in media was added to the cells for 2 hours prior to the addition of CCCP to a final concentration of 10 μM. CCCP treatment was stopped after 24-hour incubation by washing cells with PBS followed by fixation in 4% PFA. Imaging was performed on the Cytell Cell Imaging System at 10X magnification. 8 fields were captured with the same ROI positions in each well.

### Image Analysis for Mitochondrial Clearance

The mitochondrial clearance screen was quantified first by segmenting every cell in each image. Cells were stained with DAPI to visualize nuclei and whole-cell segmentation was also performed in the GFP channel, based on Parkin distribution. Cells were categorized into two groups, based on mito-DsRed intensity, a matrix-targeted fluorophore which serves as a mitochondrial marker. Cells were classified as either positive or negative for mito-DsRed signal based on a fluorescence intensity cutoff determined by assessing the fluorescence intensity signal for both the negative and positive controls for the screen, DMSO pre-treated cells treated with DMSO (negative) or CCCP (positive) for 24 hours.

The average mitochondrial clearance (% of cells with low/no mito-DsRed signal) for the positive control (DMSO+24 hour CCCP) and negative control (DMSO+24 hour DMSO) wells was determined and used to calculate the Z-factor, which is a metric indicative of screening robustness^57^. Normalized MAD z-scores were calculated for each of the compounds, as implemented in other mitophagy-related screens^58^, across two independent screening replicates. Molecules with the highest z-scores had the strongest effect on mitochondrial clearance.

### mitoQC assay to assess mitochondria-to-lysosome targeting in cells

HeLa cells were seeded onto coverslips in 6-well plates. The next day, they were transfected with Flag-Parkin using lipofectamine 2000 (Invitrogen, 11668019). 24 hours later, cells were treated with 10 μM CCCP (or DMSO as a control) in combination with 0.5 mM leupeptin and 2 μM E-64 in combination with 1 μM of the indicated small molecule treatments or DMSO for 6 hours prior to fixation.

Images were acquired on the Leica SP8 microscope. All image settings were kept consistent throughout imaging for each experiment. The ImageJ Plug-In developed by *Garriga et al.* was used to analyze the percentage of mitochondria which appear red-only^59^: https://github.com/graemeball/mQC_counter. At least 40 cells were quantified per treatment group in each replicate. Cells with at least 5 red-only puncta representing mito-lysosomes were classified mitophagic, similar to the analysis described by *Allen et al*^60^.

### *In vivo* mito-QC assay to assess mitochondria-to-lysosome targeting

7-day old flies were placed into vials containing low-melt agar in combination with the indicated treatments. Following 24-hour incubation, flies were incubated in whole-fly fixation reagent containing 1% PFA and 0.1% PBST (Tween-20 diluted in PBS) overnight at 4°C. Fly brains were dissected following PBS wash and subsequently fixed in 4% PFA for 20 minutes. After a 5-minute PBST wash and 5-minute PBS wash, fly brains were incubated in Fluoromount.

Following tissue dissection, all steps were performed protected from light. Brains were mounted onto slides and covered with #1.5 coverslips and sealed with nail polish. At least 3 independent biological trials were performed for each experiment and at least 2 fly brains were imaged for each treatment.

### Zebrafish locomotor and dopaminergic neuron assays

Zebrafish locomotion was evaluated in 96-well plate format using Zebrabox video tracking system. The Tg(*dat:EGFP*) zebrafish model and pertinent details to the quantification of dopaminergic neurons in the ventral diencephalon region of the brain has previously been described in detail^30^. For confocal microscopy of dopaminergic neurons, embryos were cleaned and housed in a 28°C incubator. 24 hours later, 1 mM MPP^+^ was administered with probucol at 50 μM concentration and DMSO control. Microscopy was performed the next day. For locomotor assays, instead of performing microscopy, media was changed to fresh MPP^+^ and drug for 2 additional days, then on the on the final day, the movement trajectory of the zebrafish was tracked using ZebraBox.

### Measuring LD and lysosome area

ImageJ was used to create maximum intensity projections of image z-stacks. The object counter Plugin was then employed to identify circular objects in the images and measure the area of puncta. Since variation in cell size was evident across experiments, the area of the cell structure of interest was normalized to the total cell area. TH staining was used to determine total area of dopaminergic neurons, while the brightfield was used to define cell area in culture.

### Statistical analysis and figures

Unpaired student’ t-tests were performed to make pairwise comparisons. Where multiple comparisons were made, ANOVA statistical analysis was performed with Dunnett’s multiple comparison correction. Independent biological replicates were used to compute statistics using GraphPad Prism v 9.0 software. Where possible, each independent biological replicate is displayed in the graphs. Error bars on graphs represent the SEM, as several technical replicates were averaged to obtain each independent replicate value. BioRender (https://www.biorender.com) was used to make Figure 1C, 6F and Figure S1A.

### Materials

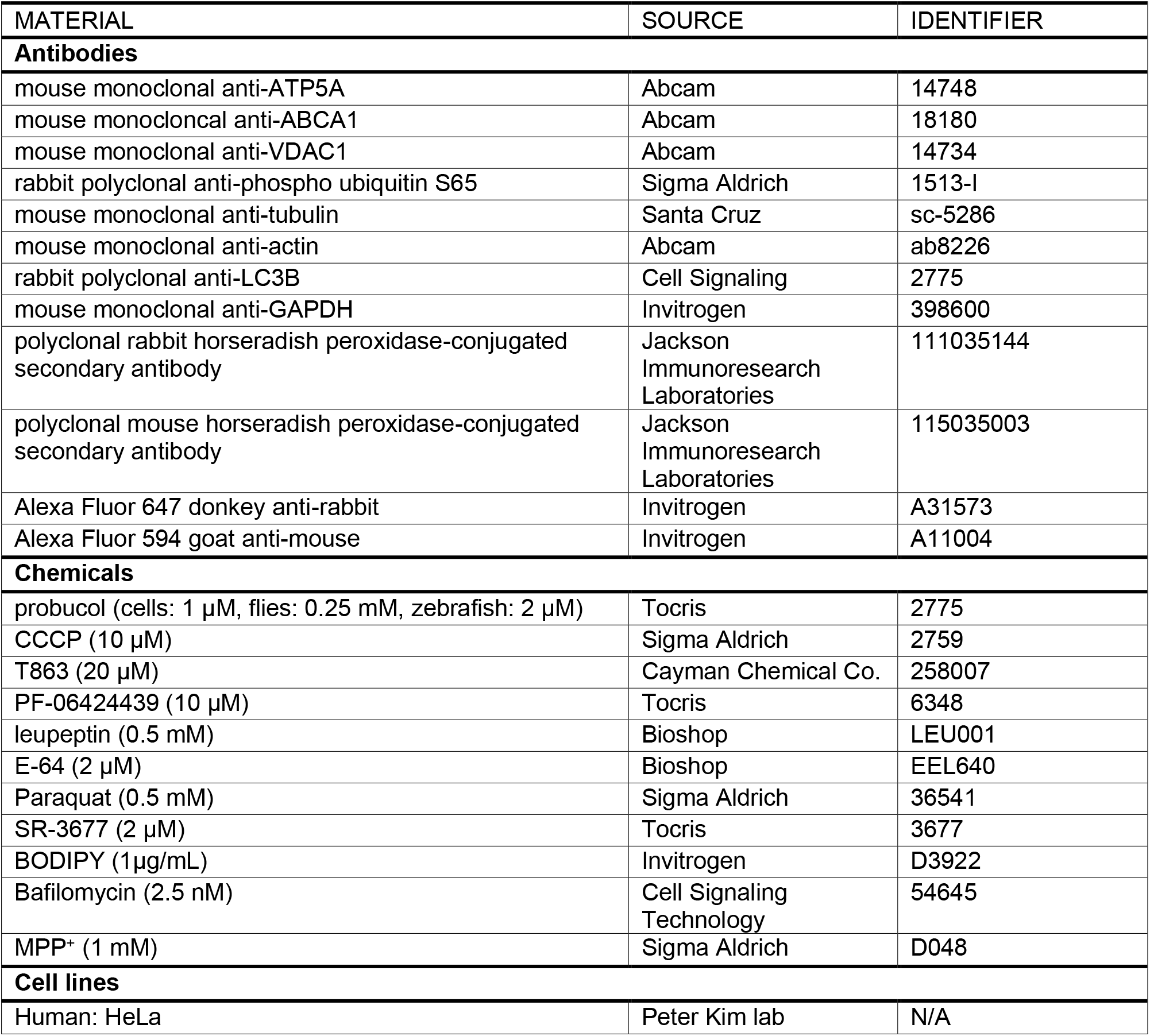
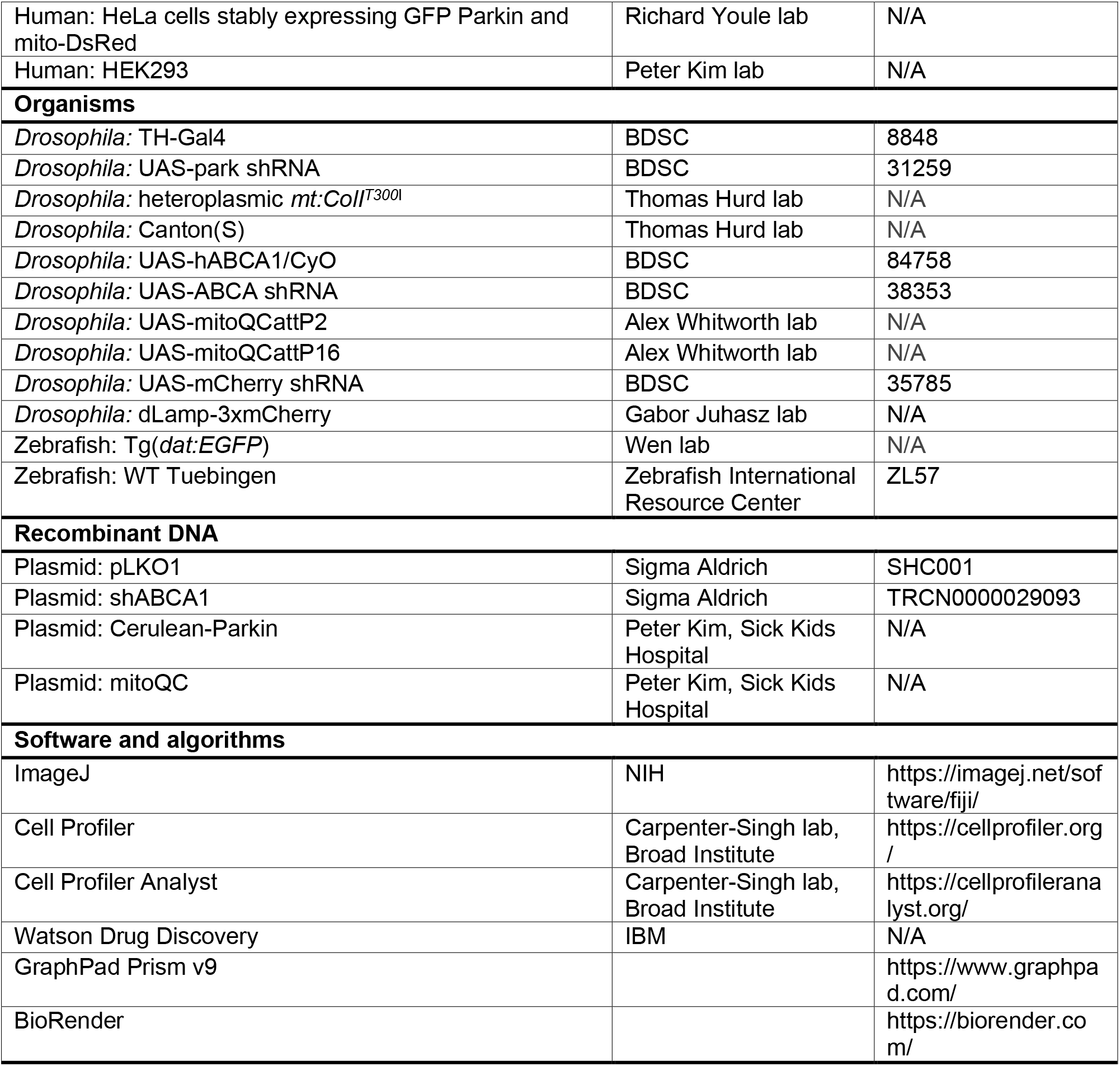

## Data availability statement

The authors confirm that the data supporting the findings of this study are available within the article and its supplementary materials.

## Acknowledgements

The authors would like to thank Dr. Guang Shi for performing microscopy for the mitochondrial clearance screen. We also thank the following labs for supplying critical tools: Dr. Richard Youle (GFP Parkin mito-DsRed HeLa cells), Dr. Thomas Hurd (heteroplasmic mt:*ColI^T300I^* flies) and Dr. Yan Wei Xi [Tg(*dat:EGFP*)].

## Appendix Table of Contents

Appendix Figure S1: Mitochondrial clearance assay screening workflow

Appendix Figure S2: Effect of screening hits on ATP5A levels following mitochondrial depolarization

Appendix Figure S3: PINK1-mediated phosphorylation of mitochondrial ubiquitin and Parkin recruitment are not affected by probucol treatment

Appendix Figure S4: Effects of ABCA1 manipulations on mitophagy

Appendix Figure S5: Lipid droplet area expansion following CCCP treatment is reduced by probucol treatment

Appendix Figure S6: Probucol improves climbing defects and survival decline which arise from mtDNA mutation *mt:ColI^T300I^*

Appendix Figure S7: Effect of probucol on LC3 lipidation under basal conditions, following mitochondrial depolarization and starvation.

Data S1: *in silico* candidates from DrugBank ranked in order of similarity to mitophagy enhancer training set

Data S2: *in silico* candidate molecules associated with terms ‘mitochondrial damage’ and ‘apoptosis’

Data S3: Results of cell-based mitochondrial clearance screen

Data S3: source data

---

**Appendix Figure S1:**
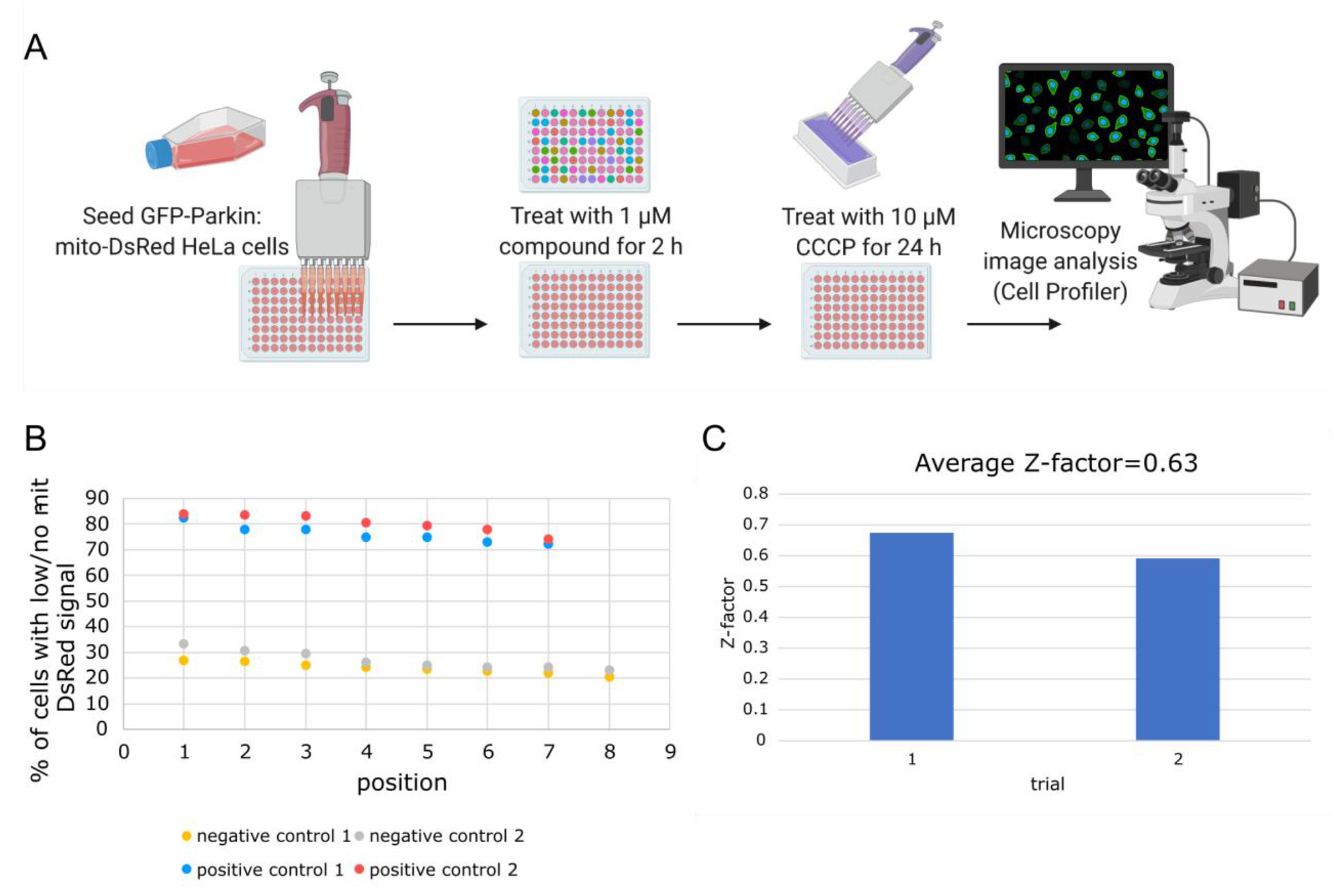
Mitochondrial clearance screening workflow. (**A**) On Day 1, GFP-Parkin mito-DsRed HeLa cells are seeded in 96-well plates. On Day 2 cells are pre-treated with 1 μM concentration of the small molecule library for 2 hours prior to the addition of 10 μM CCCP for 24 hours. Cells are then fixed and DAPI staining is performed to visualize nuclei. Cell Profiler and Cell Profiler Analyst tools are used to differentiate cells which retain mito-DsRed signal and ones with no/low mito-DsRed signal. (**B**) Mitochondrial clearance values for positive control wells pre-treated with DMSO for 2 hours in place of small molecules and followed by 24-hour treatment with CCCP to induce mitophagy and negative control wells containing cells treated with DMSO alone. (**C**) Z-factor values calculated from the positive and negative control replicate wells in each of the independent biological screening replicates.

**Appendix Figure S2:**
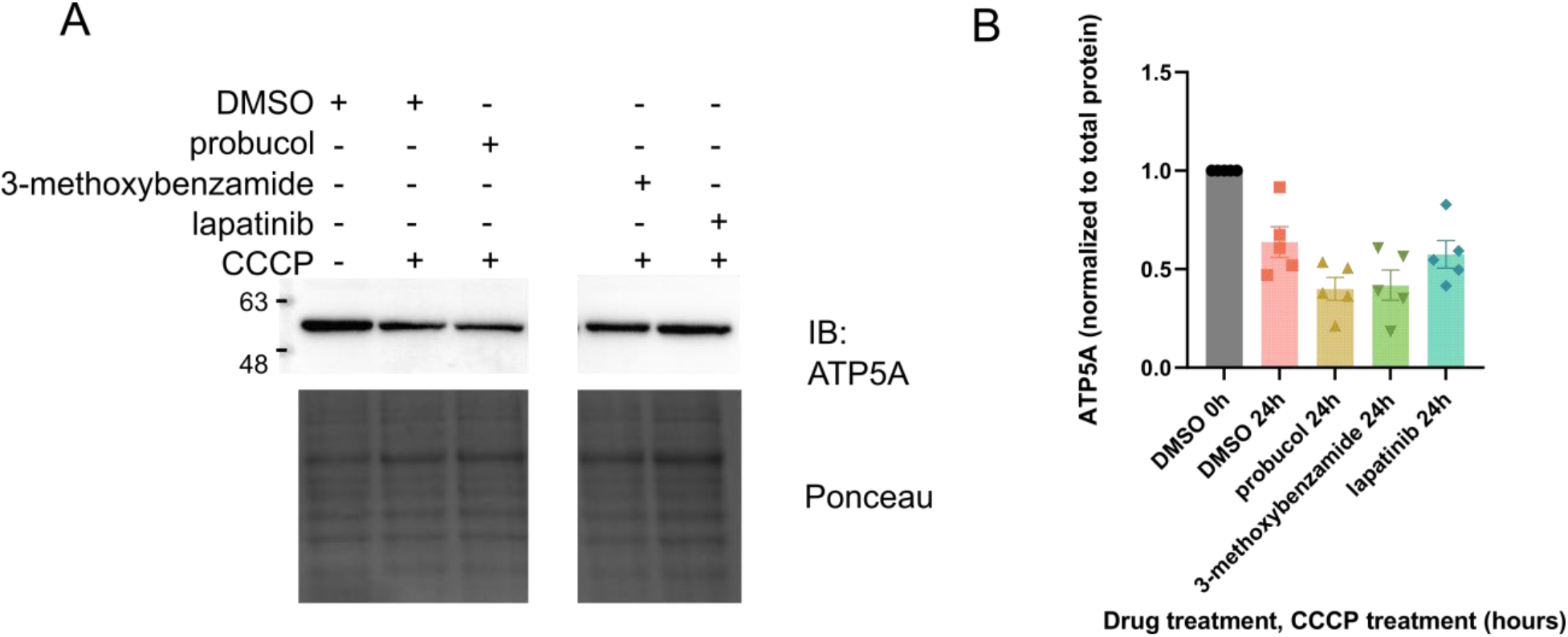
Immunoblotting to assess ATP5A levels following prolonged mitochondrial damage. (**A**) Immunoblotting whole cell lysates using antibodies for inner mitochondrial membrane protein ATP5A. HeLa cells stably expressing GFP-Parkin were treated with the indicated drugs at 1 μM concentration in combination with 10 μM CCCP. Irrelevant lane in the center of the blot was removed for clarity, but both right and left side of blot and Ponceau correspond to the same image from the same membrane. Ponceau staining was used to visualize protein loading. (**B**) Densitometry analysis was performed to assess ATP5A levels normalized to Ponceau loading.

**Appendix Figure S3:**
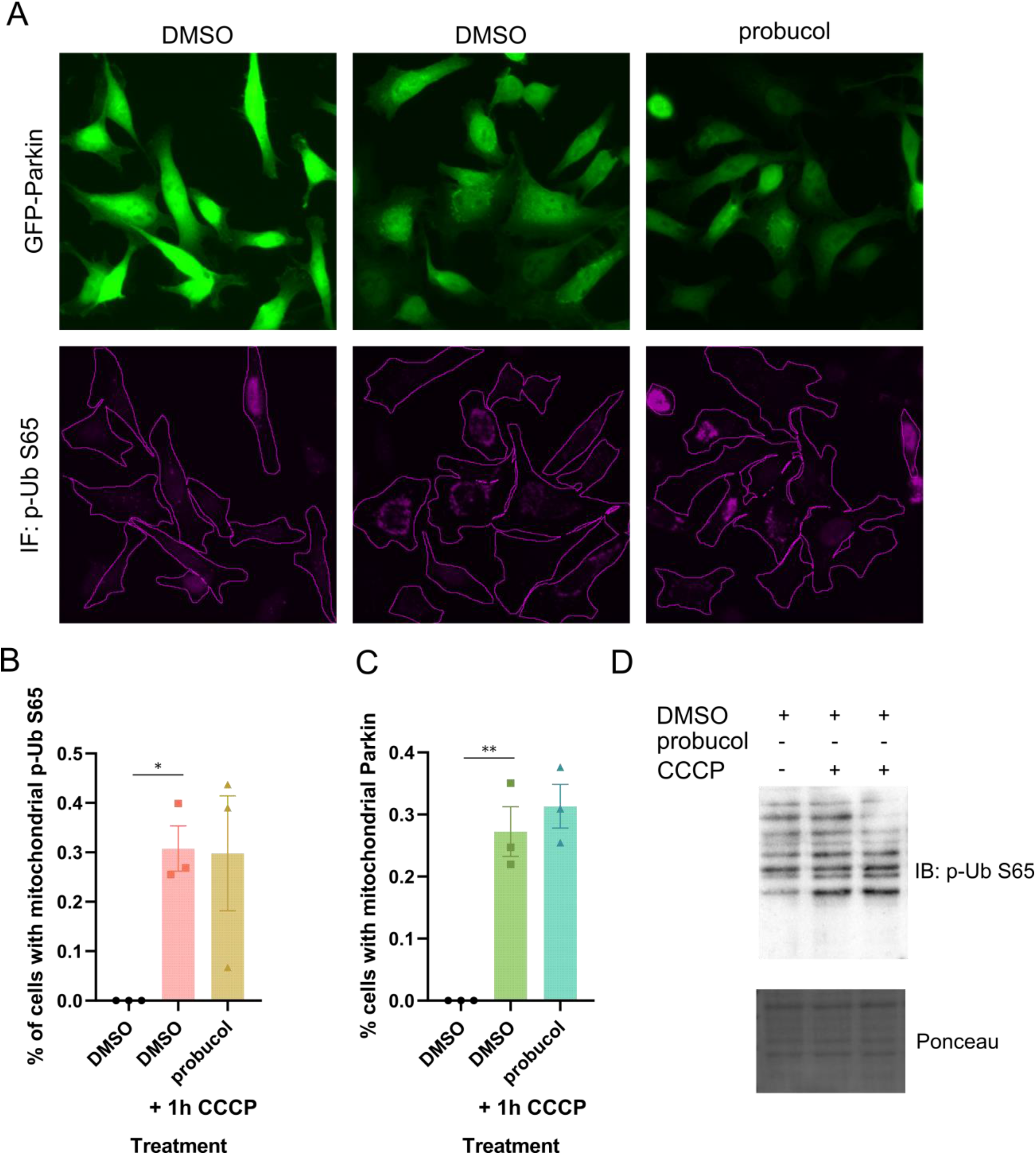
PINK1-mediated phosphorylation of mitochondrial ubiquitin and Parkin recruitment are not affected by probucol treatment. (**A**) HeLa cells expressing GFP-Parkin were pre-treated with probucol or DMSO for 2 hours prior to 1 hour treatment with CCCP. Immunostaining with antibody specific for phospho-Ubiquitin S65 (p-Ub S65) was performed on cells. (**B**) The percentage of cells which are positive for mitochondrial p-Ub S65 signal and in which (**C**) Parkin distribution is mitochondrial was calculated for each treatment. (**D**) Whole cell lysates from treatments as described in A) were probed with antibody against p-Ub S65. Ponceau staining was used to assess protein loading. Three independent biological replicates were performed for all experiments and are represented by data points and bars depict means. Error bars display SEM. ANOVA statistical analysis with Dunnett’s multiple comparison correction was performed, * and ** indicate p-value<0.05 and 0.01, respectively.

**Appendix Figure S4:**
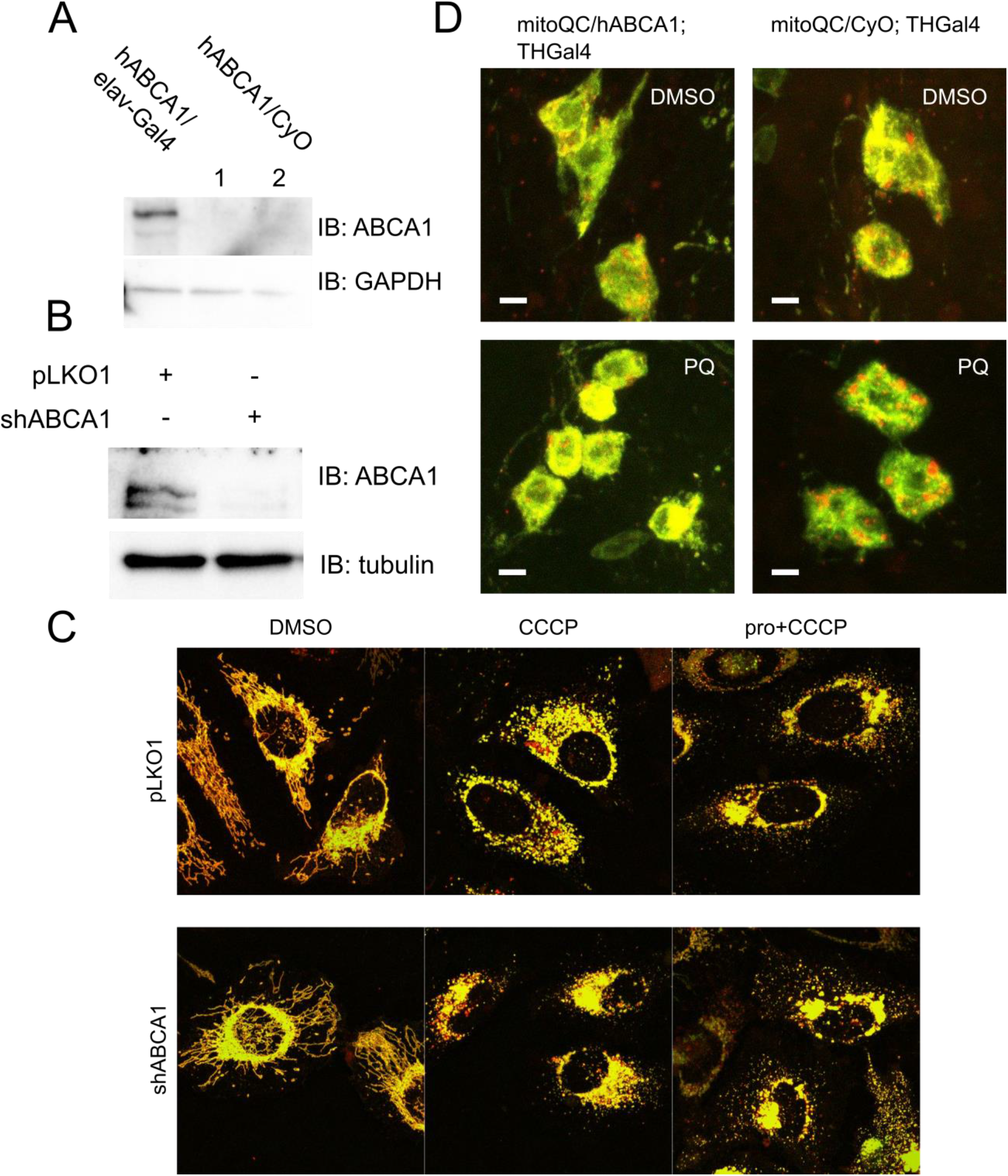
Effects of ABCA1 manipulations on mitophagy. (**A**) Immunoblotting was performed to assess ABCA1 levels in lysates derived from the brains of flies in which human ABCA1 transgene expression is driven by pan-neuronal elav-Gal4. (**B**) Immunoblotting was performed using antibodies against ABCA1 and tubulin as a loading control on HeLa cells lysates from cells transfected with either pLKO1 control or shABCA1 vectors. The blots pictured in A) and B) are representative of 3 independent biological replicates. (**C**) HeLa cells expressing the mitoQC reporter and Cerulean-Parkin were transfected with pLKO1 or shABCA1 and treated with CCCP for 6 hours. (**D**) Mitophagy was assessed in flies with expression of the mitoQC reporter in the presence and absence of the human ABCA1 transgene induced by the TH-Gal4 driver. Food containing the indicated combinations of probucol and paraquat was administered.

**Appendix Figure S5:**
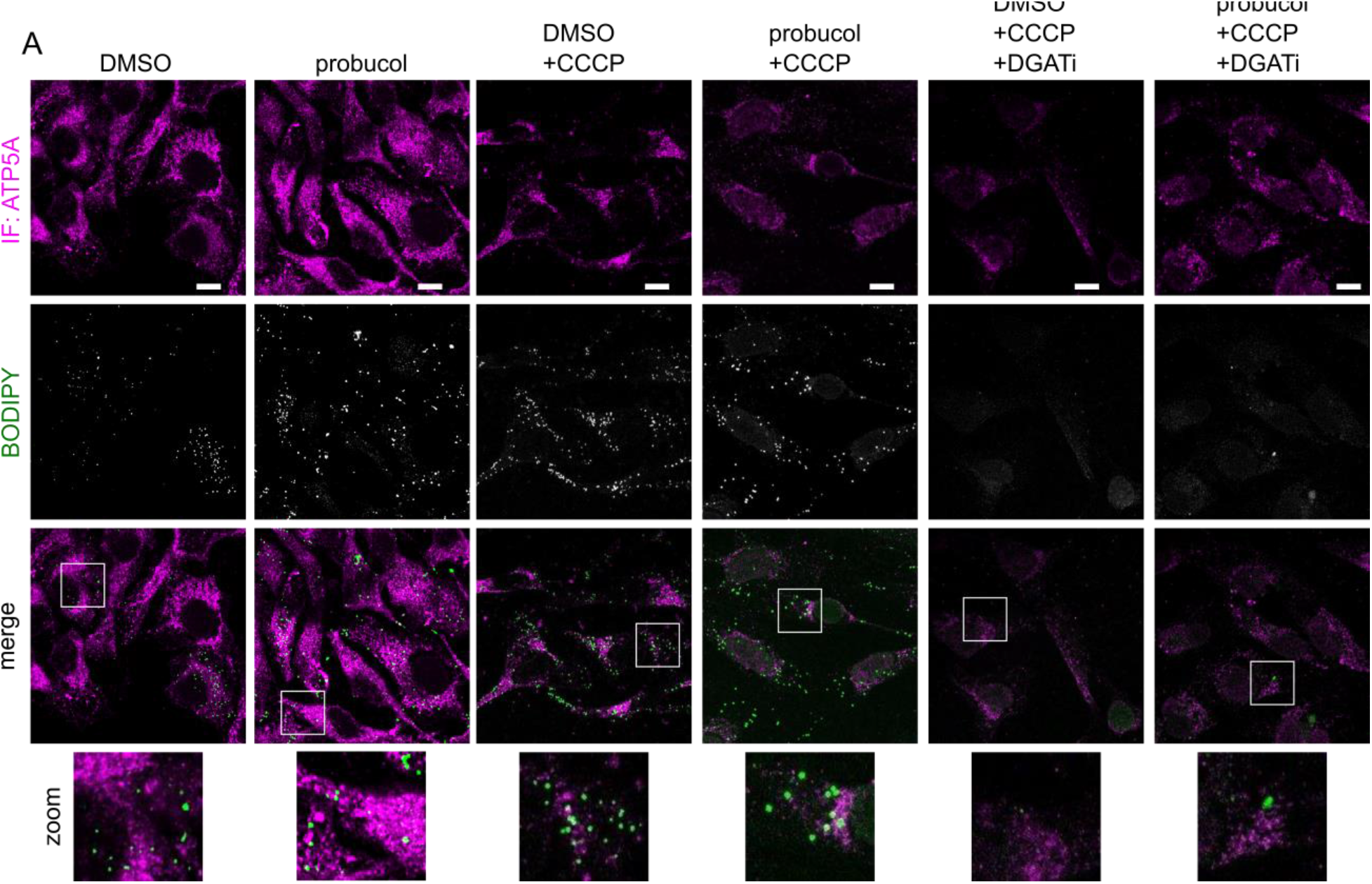
Lipid droplet area expansion following CCCP treatment is reduced by probucol treatment. HeLa cells were treated with the indicated combination of DMSO, probucol, CCCP and DGAT inhibitors. BODIPY staining was performed to visualize LDs and immunostaining with ATP5A antibody.

**Appendix Figure S6:**
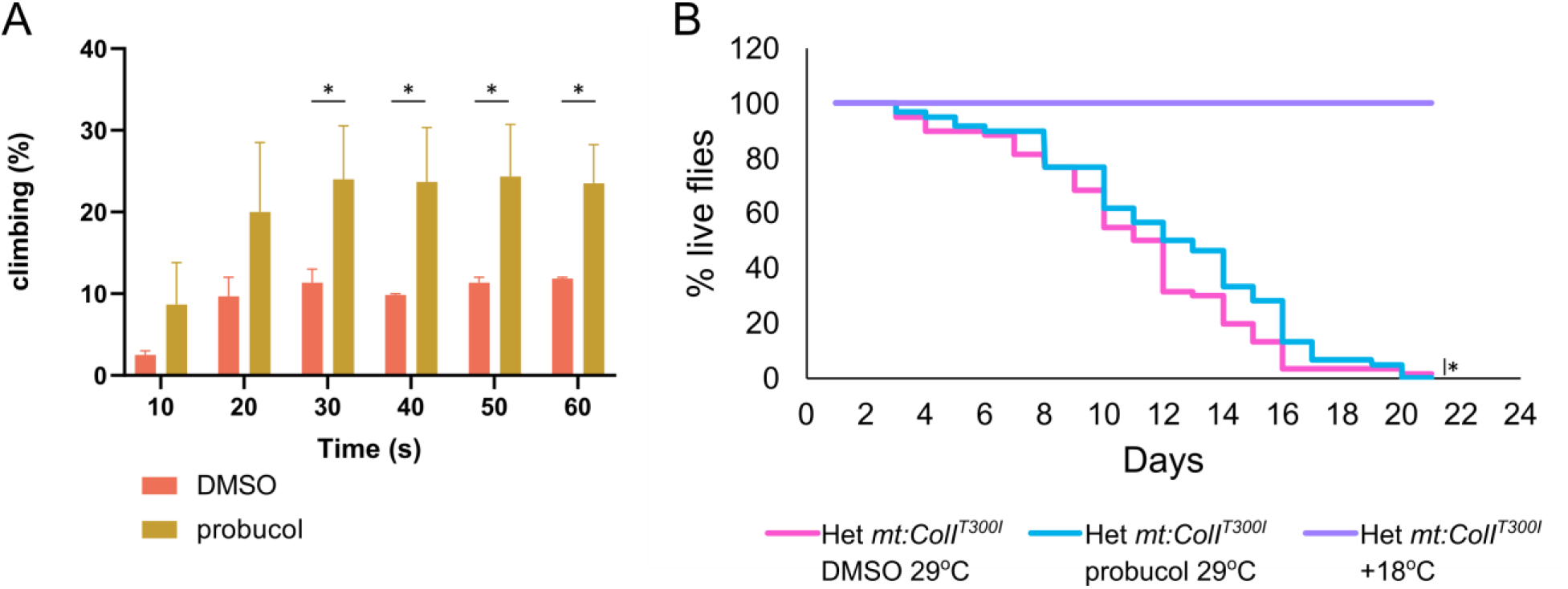
Probucol improves climbing defects and survival decline which arise from mtDNA mutation *mt:ColI^T300I^*. Heteroplasmic flies with *mt:ColI^T300I^* in approximately 90% of their mtDNA were fed food supplemented with either DMSO or probucol. (**A**) The percentage of flies climbing beyond a height of 12.5 cm is displayed, in addition to (**B**) the survival of the flies in both groups. As a control, heteroplasmic *mt:ColI^T300I^* were maintained at permissive temperature of 18°C. Three independent biological replicates were performed for both A) and B) and at least 20 flies were included in each replicate. Bars represent mean values in A and error bars represent SEM. For A), unpaired student’s t-tests were performed to evaluate differences between DMSO and probucol. For B) log-rank test analysis was performed to compare the survival of the two treatment groups housed at 29°C. * indicate p-value<0.05.

**Appendix Figure S7:**
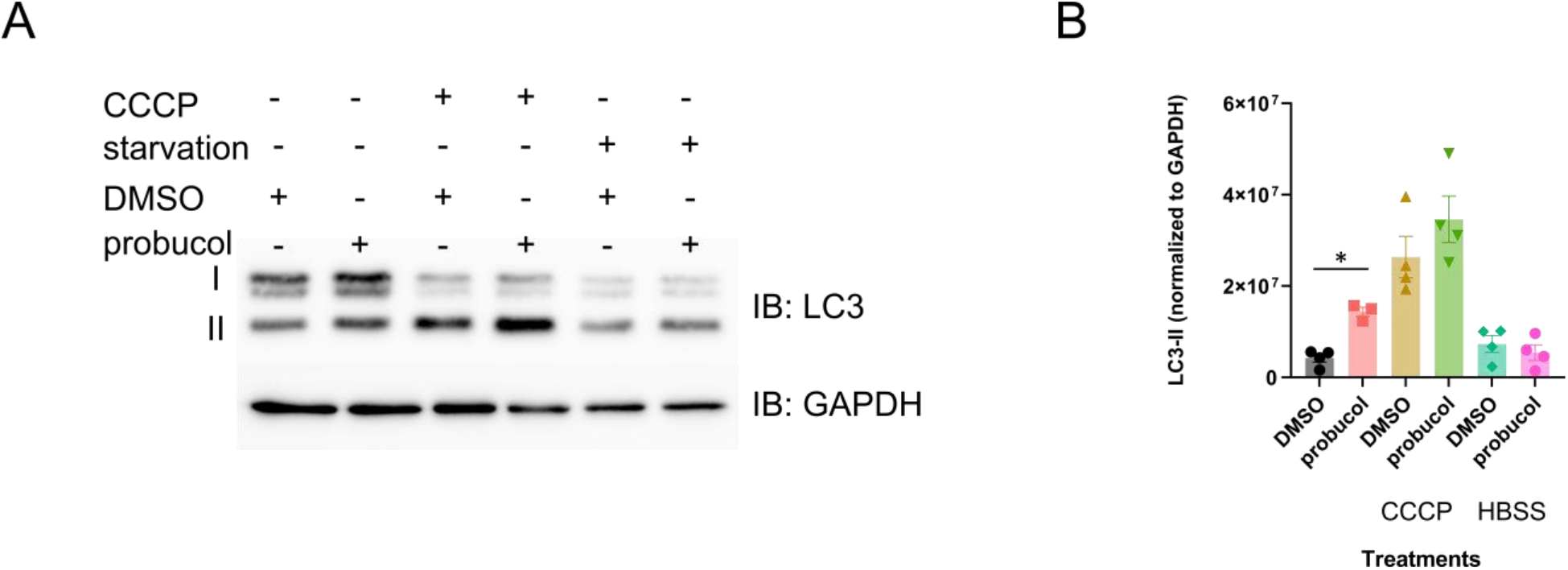
Effect of probucol on LC3 lipidation under basal conditions, following mitochondrial depolarization and starvation. **A**) HEK293 cells were either incubated in DMEM, DMEM with CCCP or HBSS media for 6 hours. Lysates were separated by SDS-PAGE and immunoblotting was performed using antibodies which recognize LC3 and GAPDH, as a loading control. **B**) Densitometry was performed to measure the levels of lipidated LC3-II, which were normalized to the GAPDH loading control. Unpaired t-tests were used to evaluate differences between DMSO and probucol. * indicate p-value<0.05.

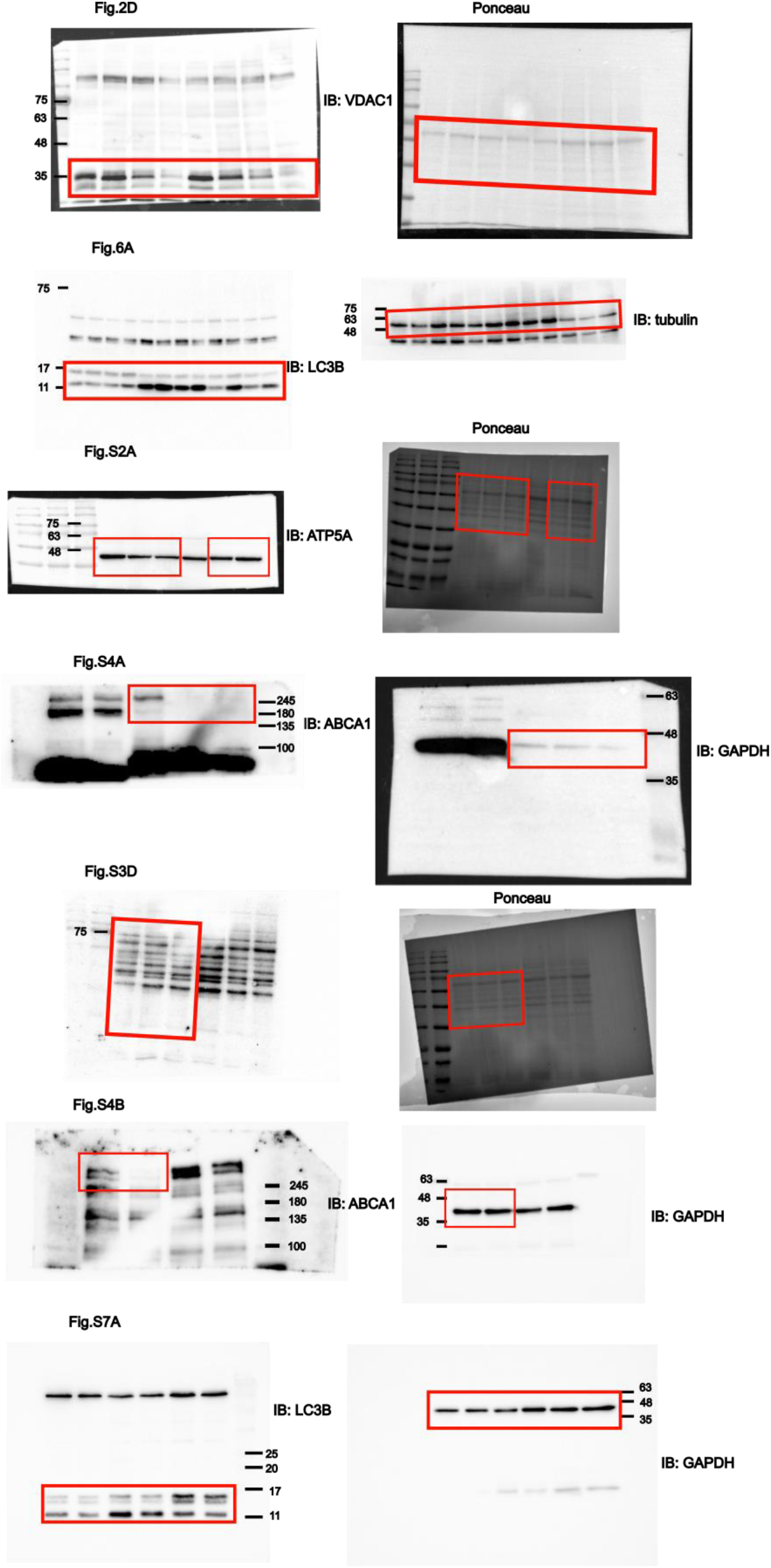

